# Control of wildtype zebrafish optomotor response with a photoswitchable drug

**DOI:** 10.64898/2026.03.05.709743

**Authors:** L. Camerin, J. Martínez Tambella, G. Schuhknecht, V. M. Wang, K. Krishnan, P. Pflitsch, F. Engert, P. Gorostiza

## Abstract

For animals to interact effectively with their environment, the brain must integrate sensory information and generate appropriate motor responses. Multiple neuronal circuits contribute to this process, and identifying their roles remains a central focus in neuroscience. The recently developed photoswitchable compound Carbadiazocine controls neuronal firing across species. It modulates larval zebrafish locomotion and alleviates neuropathic pain in rodents in a reversible, light-induced manner. Given its effects on both motor and somatosensory circuits, we investigated the impact of Carbadiazocine on sensorimotor behaviors. We focused on the optomotor response in zebrafish larvae and assessed its potential as a tool for circuit perturbation and behavioral analysis, for the first time combined with photopharmacology. We performed experiments in head-fixed and free-swimming larvae to assess their capacity to detect and follow the direction of optic flow, as well as to characterize swimming speed patterns and individual tail bout properties following administration of the two Carbadiazocine photoisomers. In both paradigms, treatment with the pre-illuminated compound led to a decrease in accuracy in responding to optic flow (correct turning percentage dropping from ∼95 % to ∼80 % in head-fixed experiments and correctness decreasing from ∼65 % to ∼20 % in free-swimming experiments). Speed analysis revealed an increased number and duration of fast movements with a decrease in number and duration of slow movements, even during periods without visual stimulation. Tail bout analysis further showed an increase in 15-30 Hz bout frequencies, corresponding to incomplete, irregular tail movements. All these effects were absent when the dark-relaxed compound was administered. Together, these findings deepen our understanding of sensorimotor transformations and lay the foundations to probe native neuronal circuits underlying behavior in diverse animal species using a wide dynamic range of photostimulation patterns.

## INTRODUCTION

Photopharmacology is an expanding field that develops spatiotemporally-controlled pharmacological compounds. Small-molecule drugs can be designed to undergo conformational changes upon light irradiation with specific wavelengths, which alter their biological effects.^[1–4]^ This isomerization process is reversible with time by relaxation back to its thermally stable state or upon irradiation with a second wavelength. The possibility of turning on and off the biological activity of chemical compounds with specific wavelengths allows to constrain their activity within tissue using illumination patterns. Preventing unwanted activity outside the region of interest thus reduces potential adverse effects.^[5–8]^ Such drug-based control of cellular activity with light can be exerted on all wildtype organisms expressing the target protein and does not require genetic manipulation. Thus, it is generally advantageous over optogenetic methods like channelrhodopsin overexpression.

The choice of suitable animal models is essential to study photoactivation inside an organism. Zebrafish (*Danio rerio*) has proven valuable to test and screen photoswitchable drugs.^[8–16]^ Zebrafish are freshwater tropical fish in the Cyprinidae family. They share approximately 70 % of their genome with humans and 84 % of human disease-associated genes have zebrafish homologs.^[10,17,18]^ Its small size and rapid development make it a promising model for studying human disease, neuronal functions, and development.^[19–24]^ The fact that larvae are naturally translucent for the first few weeks of life facilitates observation of organs and internal structures under the microscope in living organisms and in real time, as well as the capability for modulating the activity of photoswitchable drugs. This makes zebrafish larvae a versatile model for a wide variety of photopharmacological studies. Larvae have been used for toxicity and efficacy studies with several readouts, including survival rate, developmental abnormalities, heart rate modulation, tumor growth control, and visual acuity.^[10,24–26]^ Swimming activity is generally used to evaluate not only the safety profile of drug candidates, but also to assess their effects on targets such as dopaminergic or glycinergic neuronal circuits that regulate locomotion.^[13,15,27–29]^ However, these assays are often limited to quantifying whether a drug increases or decreases locomotion, and complementary experiments are required to elucidate the drug action site(s) in the neural pathway and whether other neural processes are being modulated. Among the wide repertoire of behavioral tests that can be implemented on zebrafish larvae, those combining complex sensory stimuli with specific locomotor readouts offer an added value to the standard swim distance quantification as they allow us to probe the whole sensorimotor process underlying the observed behavioral response.^[24,30,31]^ Moreover, the development of sophisticated tracking software for larval zebrafish has permitted the study of locomotor output with enough granularity to classify individual swim episodes according to their speed and duration and to study motor responses at single bout resolution.^[32,33]^

For these reasons, the zebrafish optomotor response (OMR) behavioral assay has been an instrumental tool to study complex sensorimotor processing and its underlying circuits.^[34–38]^ Fish have an innate ability to detect visual motion and follow the direction of the perceived movement, which is a critical survival mechanism that is thought to help them avoid involuntary displacement in water currents and to evade predators.^[39,40]^ The OMR in zebrafish is characterized by discrete swim bouts, each comprising a few tail oscillations, which can be analyzed as individual decisions resulting from sensory inputs. These bouts are interspersed with quiescent periods, known as interbouts, during which sensory information may be accumulated.^[32,41–45]^

The OMR in zebrafish larvae has been widely reproduced with a stimulus pattern made of projected high-contrast parallel stripes that move perpendicular to the stripe direction in the fish arena.^[46]^ In addition, the random dot kinematogram (RDK), a visual stimulus originally developed for primate studies, has been successfully adapted for zebrafish larvae.^[32,45]^ It consists of projected dots moving perpendicular to the fish body axis. When exposed to whole-field motion drift from the dots, zebrafish exhibit the OMR and move in the same direction as the dots. This behavior requires no training and is present in larvae as early as 5 days post-fertilization (dpf), making it an ideal system for exploring the neural basis of sensorimotor integration and decision-making.

OMR experiments have been combined with optogenetic control of genetically modified neurons in zebrafish, allowing to map certain circuits and to investigate sensorimotor transformations.^[34]^ However, despite the unique ability of photoswitchable drugs to control intact neurons in wildtype fish with pharmacological and spatiotemporal selectivity, OMR assays have never been performed under photopharmacological control.

Carbadiazocine is a photoswitchable analog of carbamazepine, a voltage-gated sodium channel blocker that controls neuronal firing. Upon adding dark-relaxed carbadiazocine to water, the locomotion of wildtype zebrafish larvae can be increased with light of 405 nm.^[15]^ Here, we have combined OMR assays in freely moving and head-fixed wildtype zebrafish larvae with the use of carbadiazocine in its two forms. We observe that the illuminated compound decreases the accuracy of visual stimuli responses, enhances fast swimming movements over slow ones, and causes irregular tail movements. These findings lay the foundations to probe intact neuronal circuits underlying sensorimotor transformations in diverse animal species using patterns of photostimulation at wide spatiotemporal scales.

## RESULTS

To evaluate the effect of Carbadiazocine on zebrafish behavior involving complex sensorimotor integration, we performed OMR tests with free-swimming larvae in 10 cm-long linear wells using a Viewpoint Zebrabox videotracking instrument and Zebralab analysis software (see Figure 1 and SI for details). Larvae were first exposed to complete darkness for 5 minutes, followed by 5 cycles of visual stimuli. The stimuli consisted of 10 alternating black (dark) and blue stripes perpendicular to the linear well that moved back and forth along the well at 10 rpm for 30 seconds. To verify that the blue light projected for the visual stimuli was not photoswitching the compound, controls were run with the Carbadiazocine solutions (dark-inactive and active 400 nm light pre-illuminated), see SI Figure S10.

**Figure 1.**
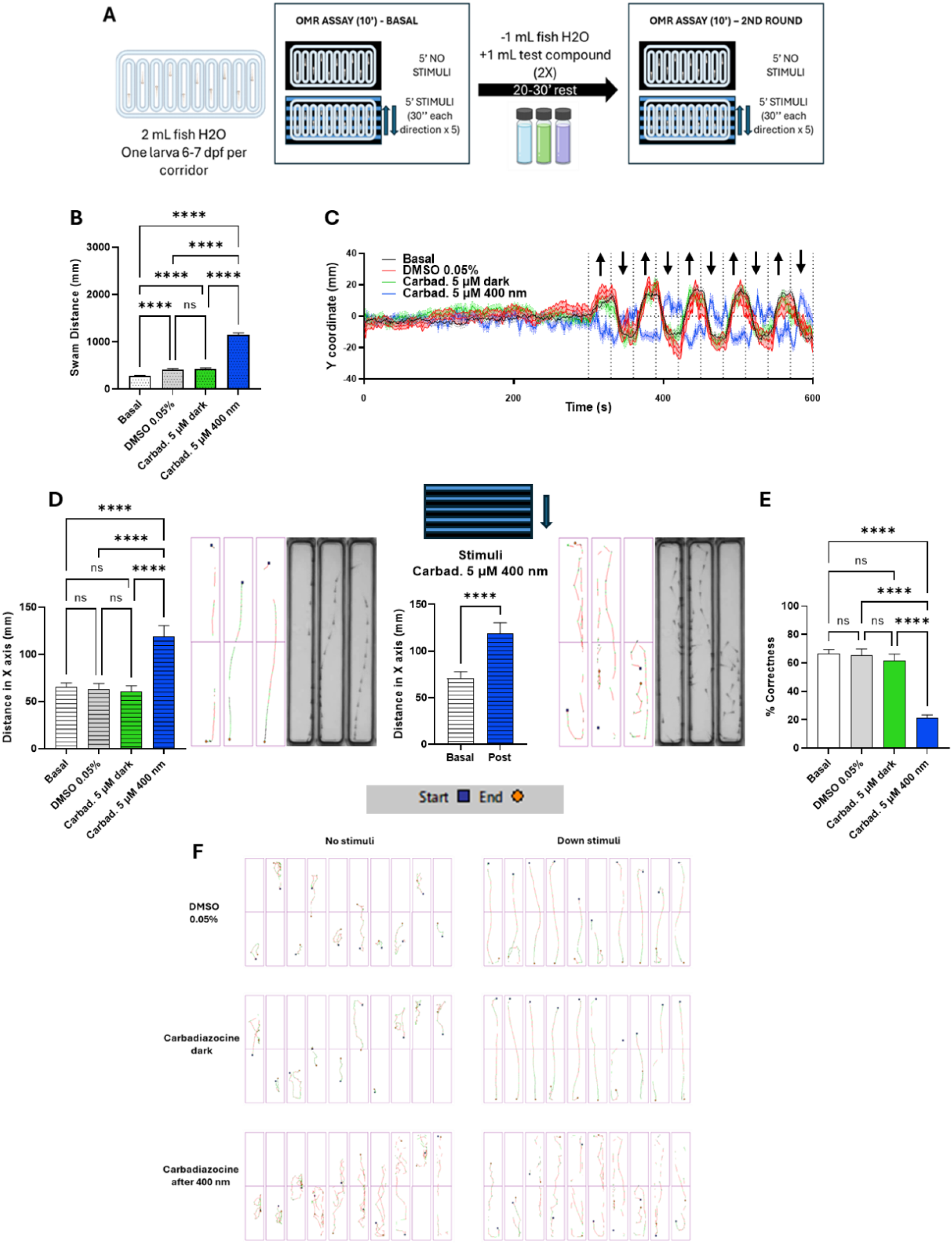
Optomotor response assay in free swimming zebrafish larvae (n=40 for DMSO 0.05 %, n=50 for benchtop and pre-illuminated Carbadiazocine). **A)** Experimental design showing 10 linear corridors for individual larvae (vertical in the figure) over a screen that projects patterns of horizontal blue and black (dark) stripes that move in the vertical direction, triggering the optomotor reflex in untreated larvae to follow them. **B)** Total distance swam by the zebrafish larvae in the protocol period without stimuli. Carbadiazocine 5 µM preilluminated with 400 nm light significantly increased the swam distance. **** p<0.0001 Mixed effect analysis with Tukey’s multiple comparisons test. **C)** Temporal trace of the vertical (Y) position of the larvae during the experiment. The horizontal dotted line shows the central position of the plate, and vertical dotted lines indicate the change in the visual stimuli. Arrows indicate the direction of the moving stripe stimuli in each period. **D)** Quantification of distance swam orthogonal to the stimuli (horizontal, X) and example of traces of the same three fish before and after the application of pre-illuminated carbadiazocine. Red lines correspond to fast movements (distance travelled at ≥ 6 mm·s^-1^); green lines correspond to intermediate movements (distance travelled at 2–6 mm·s^-1^); black lines correspond to slow movements (distance travelled at ≤ 2 mm·s^-1^). **** p<0.0001 Mixed effect analysis with Tukey’s multiple comparisons test. **E)** Percentage of correctness of the OMR assay. **** p<0.0001 Mixed effect analysis with Tukey’s multiple comparisons test. **F)** 30-second trajectories of zebrafish larvae swimming in the absence or presence of stripes stimuli moving downwards (left and right panels respectively) under different treatments (vehicle DMSO 0.05 %, dark-relaxed carbadiazocine and pre-illuminated carbadiazocine). Blue squares represent larva position at the beginning of the period and yellow circles their final position.

**Figure 2.**
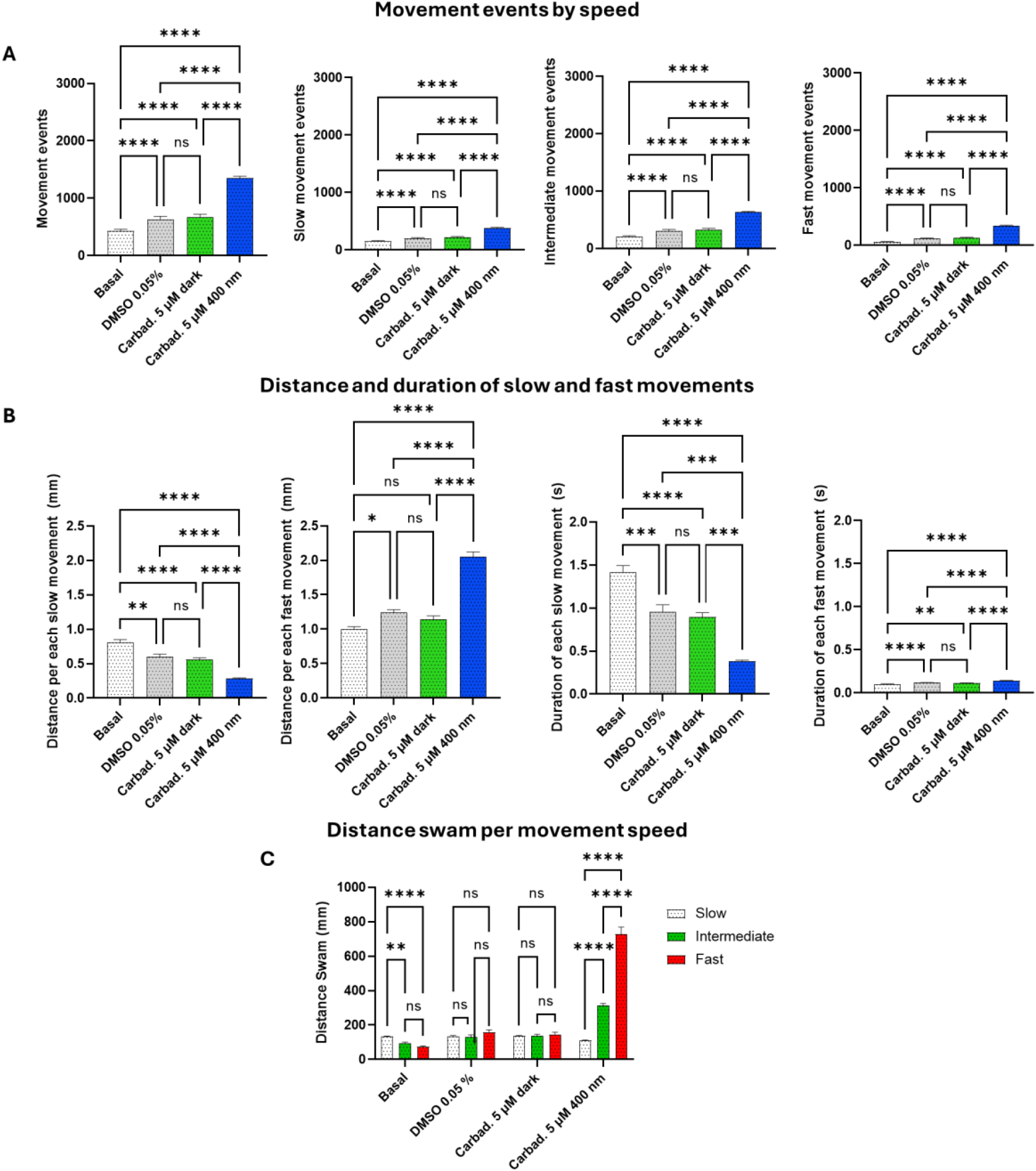
Analysis of the speed of movements during no-stimuli period of the Optomotor response assay in free swimming larvae. **A)** Count of total movement events (i.e. every time a larva initiates a movement) and classified by speed of movement for larvae in basal condition, treated with vehicle (DMSO 0.05 %, n=40), benchtop Carbadiazocine, and 400 nm pre-illuminated Carbadiazocine (n=50/group, ****p<0.0001; ns = not significant, Mixed-effect analysis followed by Tukey’s multiple comparisons test). **B)** Distance travelled and duration of each slow and fast movement for larvae in basal condition, treated with vehicle (DMSO 0.05 %), benchtop Carbadiazocine, and 400 nm pre-illuminated Carbadiazocine. ****p<0.0001; ***p<0.001; **p<0.01; *p<0.05; ns= not significant, Mixed-effect analysis followed by Tukey’s multiple comparisons test. **C)** Distance swam with slow, intermediate and fast movements for larvae in basal condition, treated with vehicle (DMSO 0.05 %, n=40), benchtop Carbadiazocine and 400 nm pre-illuminated Carbadiazocine (n=50/group). ****p<0.0001; ns= not significant, Two-way ANOVA followed by Tukey’s multiple comparisons test.

The experiment was designed in two consecutive parts (see Figure 1.A): (1) one initial phase of the recording allowed to quantify the larvae baseline swimming response to the OMR stimuli in fish water (basal); (2) next, larvae from different wells were treated with either vehicle alone (0.05 % DMSO), 5 µM Carbadiazocine under dark conditions, or 5 µM Carbadiazocine pre-illuminated at 400 nm. Carbadiazocine solutions contained 0.05 % DMSO (final concentration in the well). After the addition of these solutions, the plate was returned to the Zebrabox to reassess the OMR response. The results were analyzed by comparing the larvae initial and final positions within each well during the presence of the stimuli. A response was considered correct if the larva moved in the same direction of the stimuli (i.e., if the stripes moved upward in Figure 1 and the larva’s final position was higher than its initial position). In practice, a correctness threshold for larvae position was established as the mean Y coordinate variation of untreated larvae in absence of stimuli (see SI for details).

Untreated larvae (basal) swam relatively short distances, mainly displacing along the wells in the direction of visual stimuli (Y or vertical in Figure 1.C) and displayed high correctness (Figure 1.E), consistent with zebrafish OMR.^[47]^ Although the vehicle (0.05 % DMSO) increased the swim distance in absence of stimuli, no further increase was observed with the dark-relaxed Carbadiazocine, confirming that the *cis* relaxed isoform of the drug does not alter larvae locomotion^[15]^ (Figure 1.B). In contrast, larvae treated with 400 nm pre-illuminated Carbadiazocine showed a twofold increase in swim distance (Figure 1.D) and a three-fold decrease in correctness (Figure 1.E). The effect of the active isoform of the drug was evident immediately upon administration, even before visual stimulus onset, because the larvae exhibited non-directed hyperactivity (SI Figure S12). In presence of the visual stimulus, hyperactivity persisted and the larvae continued to modify their swimming direction each time the OMR stimulus changed (up or down), albeit in the opposite direction to the stimulus in the case of pre-illuminated Carbadiazocine (Figure 1.C and SI Figure S11, movies M1 and M2). Moreover, these treated larvae presented a significant increase in the distance swam orthogonal to the stimuli (X direction, Figure 1.D). The hyperactivity observed in absence of moving stripes, along with the changes in swimming direction and behavior in response to visual stimuli, could be due to the drug interfering with a motor circuit (e.g., producing an under/overshoot). Interestingly, such interference does not impair the fish’s ability to detect the presence of visual stimuli in their environment.

The observed results regarding hyperactivity are consistent with the action of Carbadiazocine on the swimming behavior of both blinded and non-blinded zebrafish in circular wells.^[15]^ The effects of the drug isomers on the correctness of movements (time course and direction of OMR trajectories) have not been previously reported and provide new insights into its underlying circuits. In particular, they show that correctness can be controlled with pharmacological selectivity and offer the prospects of spatial and temporal selectivity by using suitable light patterns (e.g. to photoswitch carbadiazocine in selected groups of neurons, in individual ones, or even in specific subcellular regions like axon hillocks or nodes of Ranvier). This convenient manipulation parameter was not previously available in advanced behavioral studies like OMR that use naturalistic stimuli to evoke innate behaviors in wildtype individuals.

To better understand the characteristics of larvae movement in the different conditions, we quantified the speed of each movement event and classified them as fast (> 6 mm·s^-1^), intermediate (2 to 6 mm·s^-1^), and slow (< 2 mm·s^-1^). During the initial no stimuli period, pre-illuminated Carbadiazocine increased the total number of movement events across all speed categories (Figure 1.A). However, the duration and distance swam per movement were significantly reduced for slow movements and increased for fast movements (Figure 1.B), consistent with a decreased total distance travelled at speed < 2 mm·s^-1^ and a greater distance covered with movements exceeding 6 mm·s^-1^ (Figure S13 and S14). Pre-illuminated Carbadiazocine altered the distribution of distance travelled across speed categories during the no-stimuli period, shifting larval displacement from predominantly slow exploratory movements to a swimming behavior dominated by fast bursts (Figure 1.C) even in the absence of visual stimuli to guide behavior. Overall, these results suggest that Carbadiazocine may impair or prematurely interrupt slow movements —reducing their duration and distance— while increasing the frequency and time span of fast swims. These outcomes must result from different effects of drug isomers on the neural circuits controlling locomotor behaviors. The site(s), timing, and mechanism of action can likely be investigated using spatiotemporal patterns of drug photoswitching as discussed above.

The free-swimming OMR assay reveals that pre-illuminated Carbadiazocine profoundly alters larvae swimming patterns beyond a simple increase in locomotion. During the no-stimulus period under different treatments, we observed changes in movement duration, range, and frequency across different speeds. This hyperactivity disrupted the OMR response, causing larvae to perform irregular swims—including movements orthogonal to, and even against, the direction of the visual stripes— thereby impairing their ability to properly follow the visual cues.

While this assay provides an integrated measurement of overall swimming behavior throughout the experimental period, it lacks the precision to analyze the larvae’s initial responses to stimulation at individual swim bouts level. To address this limitation, we employed head-fixed experiments in which the larvae’s heads are immobilized in agarose gel while the tail remains free to move, allowing for detailed analysis of individual swim bouts. Furthermore, we replaced striped patterns by an RDK as visual stimulus, which provides a more controlled and refined stimulus for probing zebrafish larval behavior in neuroscience.^[32]^ The RDK assay was previously used by Bahl and Engert to validate experimentally the probabilistic decision-making model they developed to predict how zebrafish larvae exposed to a visual stimulus can integrate and retain sensory evidence over several seconds and provide a valuable model for investigating neural circuit and decision-making processes.^[32,45]^

The RDK-evoked OMR assay can also benefit from the ability of turning on and off selected neurons using photopharmacology, to add granularity and facilitate the search for correlations between circuits and behavioral outcomes. Therefore, we evaluated Carbadiazocine in RDK-OMR experiments using larval zebrafish at 5-7 dpf. Larvae were embedded in transparent agarose, leaving the tail free to move in response to stimuli and leaving the mouth exposed to the medium to allow drug uptake. Excess solidified agarose was removed from the dish, as the gel can serve as a water reservoir and dilute the concentration of drug solutions. The animals were exposed to the random dot motion stimuli projected on the bottom of the arena and their behavior was tracked in real time (see SI methods) with a custom-written Python 3.7-based software package. The stimulus was designed to be perpendicular to the larvae body axis and in a closed loop with larvae movements (Figure 3). Each detected swim bout was defined as correct if the first movement of the tail followed the direction of the stimulus. Increasing levels of coherence (defined as the percentage of dots moving in the same direction in the pattern) induce an increased accuracy in the decision-making process (correctness) and a decrease in the delay in responding (time to bout) in 5-7 dpf larvae.^[32]^

**Figure 3.**
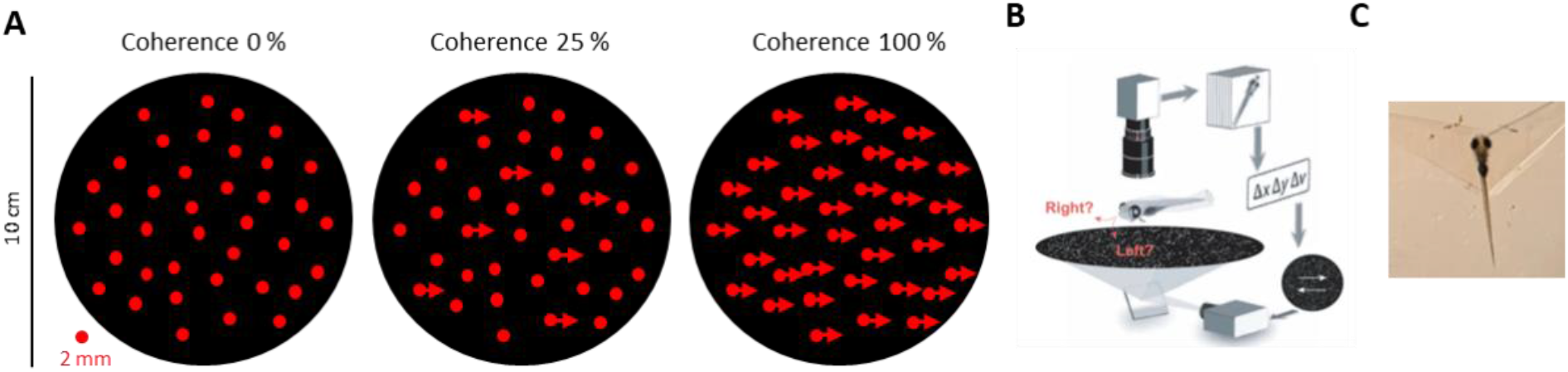
**A)** Representation of random dot kinematogram (RDK) visual stimuli at 0%, 25% and 100% coherence (e.g. rightwards) adopted from Bhal and Engert (2020).^[32]^ The real stimuli applied to the larvae include around 1,000 dots, each 2 mm in diameter, projected on a circular area with 10 cm diameter. The white light from the projector was filtered (filter cut-on wavelength 620 nm) to obtain red dots at wavelengths that are not interfering with the switching of Carbadiazocine. **B)** Representation from Bhal and Engert (2020):^[32]^ experimental set up with the visual stimuli projected from the bottom, perpendicular to the larvae. The camera and lens from the top enables tracking tail movements with a custom-written Python 3.7-based software package. **C)** Zebrafish larva embedded in agarose leaving free mouth and tail, respectively to allow drug uptake and tail movements in response to stimuli.

We conducted experiments to test Carbadiazocine, dividing the larvae into three groups for each experimental session: (1) fish treated with either 10 μM or 30 μM of the dark thermally relaxed isoform (100 % *cis*), (2) fish treated with a solution pre-illuminated with 405 nm (50 % *trans*^[15]^), and (3) vehicles (fish water with 0.1 % and 0.3 % DMSO respectively). After embedding, larvae were left to rest for at least 10 min in filtered embryo water (E3) before they were exposed to the visual stimulus for 1 h. This duration ensured sufficient trials at each coherence level, allowing to record baseline swimming activity and responsiveness to the stimuli. The fish water was then removed and changed with the prepared solutions (dark, pre-illuminated compound or vehicle respectively). After an incubation period of 20 min, the larvae underwent another hour of exposure to the random dots motion.

The swimming activity of each fish in head-fixed experiments was recorded and represented as raw traces of individual larvae tail movements (see representative examples in SI Figure S8). The traces show all tail movements, regardless of the threshold for automated bout detection and regardless of the visual stimuli. Larvae treated with either the vehicle or the dark-relaxed compound showed no change in swimming activity before and after drug administration. However, fish treated with 405 nm pre-illuminated Carbadiazocine exhibited distinct behavior compared to controls: following drug addition, the frequency of subthreshold small swims increased, interspersed with periods of complete inactivity (SI Figure S8 and Figure 4.A). The increased swimming activity agrees with the increased locomotion of blinded zebrafish larvae treated with Carbadiazocine at 400 nm.^[15]^ The inactivity periods may be due to exhaustion following hyperactivity.

**Figure 4.**
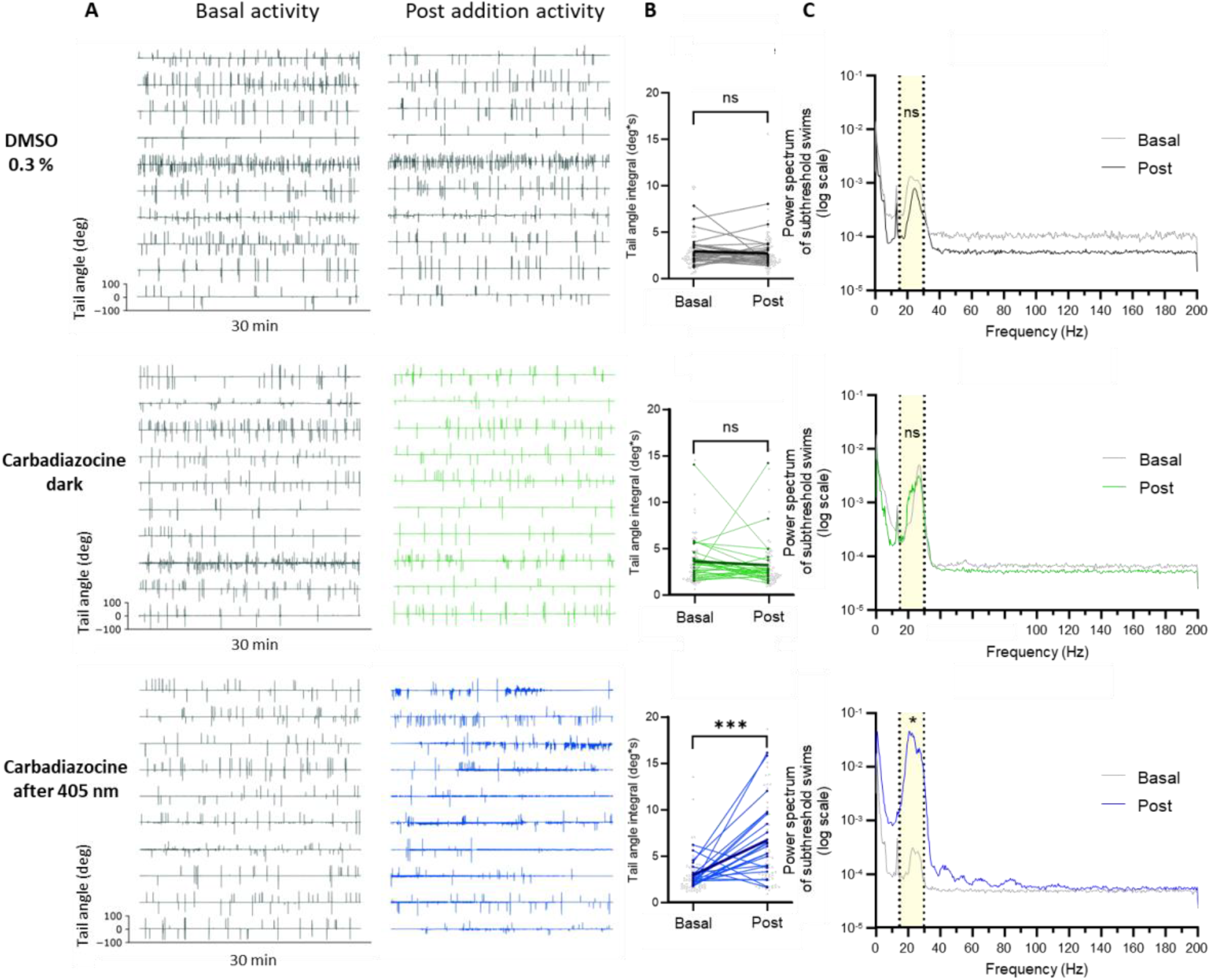
**A)** Raw traces of tail movements from individual fish illustrating basal swimming activity in fish water and swimming activity post addition of the respective test solutions (vehicle 0.3 % DMSO, dark Carbadiazocine, and 405 nm pre-illuminated Carbadiazocine). **B)** Computed tail-angle of subthreshold swims (0 % coherence trials) by integrating the absolute value of the baseline-corrected traces. Wilcoxon test, *** p<0.0003. Control 0.3 % DMSO: n=30; Carbadiazocine. Dark: n=28, data point cut off (d.p.c.o.) = 1; Carbadiazocine. 405 nm: n=23, data point cut off (d.p.c.o.) = 3. **C)** Frequency components and power spectrum of 0 % coherence traces (subthreshold swims). The area under the curve of the band between 15-30 Hz was quantified and normalized to the total area under the curve (see SI for quantification). ns = not significant; *p<0.05 post vs basal, Wilcoxon matched pairs signed rank test.

The interbout interval (time between bouts) is not a reliable parameter to quantify in our setup because the tracking software only detects as bouts the tail movements exceeding a manually pre-set threshold (see SI Methods). Small subthreshold swims induced by treatment with 405 nm pre-illuminated Carbadiazocine were not detected by the tracking software and could only be observed and quantified through raw traces of the larvae’s swimming activity. Setting an appropriate bout detection threshold was crucial to ensure that small swims did not interfere with the initiation of new visual stimulus trials (see SI Methods for details on the visual stimulus). Therefore, to study the increase in the subthreshold swims that are not considered bouts by the software, we measured the tail angle integral for 0 % coherence trials, which showed a significant increase only upon application of pre-illuminated Carbadiazocine (Figure 4.B). Moreover, the power spectrum of these subthreshold swims was analyzed, evidencing that pre-illuminated Carbadiazocine induced an increase in the proportion of swims with a tail movement frequency between 15 and 30 Hz (Figure 4.C). The increased swimming frequency observed in traces and videos of head-fixed larvae (see SI movies M3 and M4) agrees with the increased free locomotion observed in blinded larvae in 405 nm-illuminated Carbadiazocine.^[15]^

Using this setup and analysis method, we aimed to investigate whether Carbadiazocine interfered with the well-established ability of zebrafish larvae to integrate motion-derived sensory evidence and produce robust decision-making behavior.^[32,45]^ Experiments with 10 μM Carbadiazocine only showed significant difference for the pre-illuminated isomer at 100 % coherence (see SI Figure S2), possibly due to the agarose absorbing the drug and diluting it in the medium. Therefore, only the results from experiments at 30 μM were analyzed. Three coherence levels were used (0-25-100 %) with higher values yielding increased probability for correct bouts.^[32]^ We analyzed the correctness of the bouts based on the first tail deflection, establishing a threshold for bout detection (see SI Methods). We observed that in larvae treated with vehicle (0.3 % DMSO) or dark-relaxed Carbadiazocine (100 % *cis*), correctness increased with the coherence level, as reported for untreated larvae (Figure 5).^[32]^ In contrast, larvae treated with 405 nm pre-illuminated Carbadiazocine (50 % *trans*) showed lower correctness in response to the visual stimuli, in agreement with observations from OMR assays. Notably, the dark-relaxed isomer of Carbadiazocine did not affect performance at any coherence level (Figure 5).

**Figure 5.**
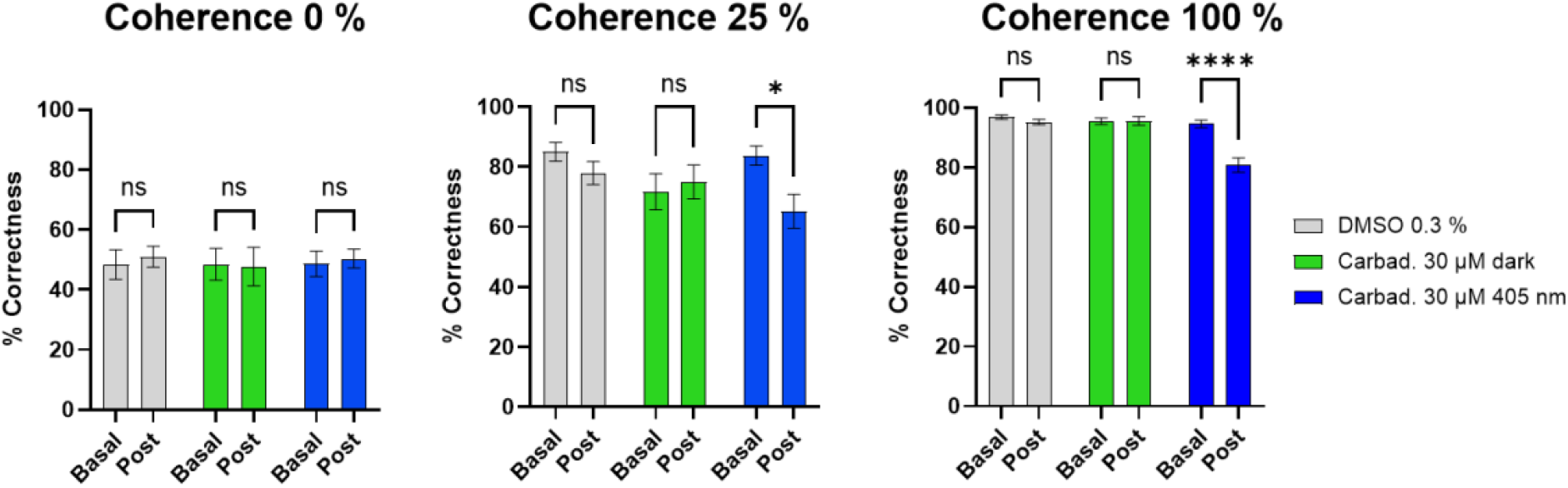
Percentage of correctness for each level of coherence (0 %, 25 %, and 100 %). Correctness drops in zebrafish larvae treated with 405 nm pre-illuminated Carbadiazocine in the presence of visual stimuli (25 % and 100 % coherence). “Basal” represents the fish performance in fish facility water; “post” indicates the fish performance after addition of the prepared solutions (vehicle 0.3 % DMSO, dark Carbadiazocine at 30 μM and 405 nm pre-illuminated Carbadiazocine at 30 μM). **** p<0.0001 Two-Way ANOVA with Šídák’s multiple comparisons test. Error bars represent S.E.M.

Taken together, these results indicate that Carbadiazocine does not alter movement correctness under dark conditions compared to vehicle controls (0.3 % DMSO) but selectively impairs performance following irradiation at 405 nm.

## DISCUSSION

In this report we combine for the first time advanced OMR assays with photopharmacology. The two behavioral assays implemented in zebrafish larvae (head-fixed and freely swimming conditions) constitute a platform to study the photopharmacological activity of Carbadiazocine in sensorimotor processes and provide greater granularity than the conventional locomotion assay to characterize their underlying neural circuits. Together, the two assays yield a complete assessment of the visually guided behavior of zebrafish larvae, each contributing valuable and complementary data. As reported, Carbadiazocine exerts isomer-dependent effects on locomotion that have now been profiled in detail, including the type of movements affected, their number, duration, and distance, along with the bout frequency range that is altered by the drug. Remarkably, in both assays the effects of the drug are only observed after illumination, whereas the minor changes observed with the dark-relaxed isomer are due to the vehicle (0.05 % DMSO).

Interpreting these results requires considering the complex neural architecture of the OMR. The OMR is a multi-step visually guided behavior that engages a distributed neural circuit: visual motion is first detected by the retina and relayed via retinal arborization fields to the pretectum and optic tectum, where directional information is processed and integrated; descending motor commands are then transmitted through premotor nuclei — including the nucleus of the medial longitudinal fasciculus (nMLF) and reticulospinal neurons, with potential cerebellar modulation — to spinal cord circuits that drive tail musculature.^[34]^ Critically, this circuit also supports the temporal accumulation of sensory evidence, a property particularly relevant when behavior is assessed over extended timescales.^[32]^ Molecular tools to selectively control each of these stages of the circuit can provide further insights into their roles and contribution to the integrated sensory-motor response.

Interpreting these results requires considering the complex neural architecture of the OMR. The OMR is a multi-step visually guided behavior that engages a distributed neural circuit: visual motion is first detected by the retina and relayed via retinal arborization fields to the pretectum and optic tectum, where directional information is processed and integrated; descending motor commands are then transmitted through premotor nuclei — including the nucleus of the medial longitudinal fasciculus (nMLF) and reticulospinal neurons, with potential cerebellar modulation — to spinal cord circuits that drive tail musculature.^[34]^ Critically, this circuit also supports the temporal accumulation of sensory evidence, a property particularly relevant when behavior is assessed over extended timescales.^[32]^

Zebrafish OMR performance is affected to a different extent in the two visually guided behavioral assays. Correctness decreases more sharply in the freely swimming paradigm (from ∼65 % to ∼20 %) than in the head-fixed paradigm (from ∼95 % to ∼80 %). This result likely arises from differences in temporal integration and movement-analysis resolution between the two approaches. While the head-fixed assay specifically evaluates whether the initial swim bout (ms scale) is aligned with the direction of dot motion, the freely swimming assay integrates locomotor responses over a 30-second stimulation period.

The relatively preserved correctness in the head-fixed condition (80 % in Figure 5) suggests that the initial perception and directional decision based on sensory input remain largely intact following treatment with pre-illuminated Carbadiazocine. In contrast, when behavior is assessed over longer time scales in freely swimming larvae, the compound-induced hyperactivity appears to impair sustained OMR performance — as if the animals skidded or overshot. Although many treated larvae can still initiate movement in the correct direction (Figure 1F), their subsequent trajectories often become irregular, with swimming bouts that are orthogonal to or even opposite the direction of optic flow (Figure 1CD). As a result, the accumulated ΔY displacement over the entire stimulus period no longer reflects proper tracking of the moving pattern by the larvae.

Consistently, changes in the average Y coordinate upon stimulus presentation indicate that treated larvae can detect the visual stimulus but fail to follow it reliably and continuously (Figure 1C). The lower correctness observed in the freely swimming assay may therefore reflect the accumulation of these irregular movements over time, amplifying subtle initial impairments into pronounced performance deficits. To further clarify the effect of Carbadiazocine in visual perception, complementary behavioral paradigms and additional models, such as blinded larvae, will be required in future work.

Several parameters can be recorded and quantified in head fixed larvae experiments (time to bout, number and fraction of completed trials, percentage of correctness over time, etc; see SI). Although most of them did not display significant changes with Carbadiazocine isomers, these metrics can reveal insightful effects of other photoswitchable and non-photoswitchable drugs, like future analogs of Carbadiazocine or other photoswitchable ligands with improved photopharmacological properties. The combination of OMR assays and a batch of diverse photoswitchable drugs targeting several neuronal proteins may thus allow profiling the circuits involved in complex behaviors. Interestingly, OMR was recently used to investigate whether silencing sensory experience through chronic exposure to tricaine during early developmental stages affects the emergence of complex sensorimotor behaviors.^[48]^ While this study focused on chronic administration—demonstrating no change in larval performance following washout of the anesthetic—we instead integrated the OMR assay with acute treatment using Carbadiazocine to induce behavioral perturbations.

Besides providing a complete characterization of the zebrafish larvae locomotor modulation by Carbadiazocine and its effects on sensorimotor behavioral assays, the results presented here show great prospects for this molecule as a tool to study neural circuits underlying different types of movement and decision-making processes. In the zebrafish larvae brain, a cluster of ∼300 descending projection neurons are connected to the spinal cord.^[20]^ These neurons can be genetically labelled with fluorescent calcium and voltage sensors and studied with advanced imaging techniques. They exhibit distinct morphological and functional properties, enabling the coordination of various behavioral outputs. Systematically testing their activity upon Carbadiazocine administration combined with precise spatiotemporal activation using light, would provide deeper insights into the contribution of specific neuronal subpopulations within this cluster to the different movement types that are modulated by Carbadiazocine. Importantly, the reversibly photoswitchable properties of Carbadiazocine, together with the absence of behavioral effects of its dark-relaxed isomer, allow exerting precise activity patterns on neurons, making it a powerful tool for circuit analysis.

The spatial resolution of photostimulation depends on the illumination method (continuous wave or multiphoton)^[14,49–51]^ but it is diffraction-limited to a fraction of a micrometer for ultraviolet and visible wavelengths used in photopharmacology. It can also be limited by diffusion of the compound. In neurons, an axial resolution of ∼5 µm has been achieved with a diffusible ligand^[50]^ and ∼1 µm on the image plane with a tethered one, allowing to control the activity of individual synapses *in vivo*.^[51]^ Thus, combining spatiotemporal photostimulation patterns (by single- or multi-photon laser scanning or holography) with OMR-based assays will allow delineating neural circuits and understanding the corresponding mechanisms at the level of synaptic functional units. Note that, in contrast to optogenetic manipulations, which operate at the cellular scale by means of available gene promoters, photopharmacology offers to directly control endogenous proteins in their physiological context and thus leveraging their corresponding signaling pathways and downstream biological processes, including behavioral outcomes.

In summary, the combination of freely moving and head fixed OMR assays with photopharmacology and the prospects of using high resolution neuronal activity imaging and manipulation with photoswitchable drugs, together lay the foundations to probe intact neuronal circuits underlying sensorimotor transformations in diverse animal species and wide spatiotemporal scales.

## CONCLUSIONS

In this work, we have investigated the impact of the photoswitchable compound Carbadiazocine on sensorimotor behavior and the ability to detect and follow visual stimuli of zebrafish larvae. We have analyzed the effects of the drug isomers on non-stimulated and visually guided locomotion by combining two advanced complementary assays. They allow for the classification and quantification of the movements of the larvae in response to different visual stimuli (moving dots at different coherence levels and moving stripes) and in different configurations (freely swimming and head fixed). Each assay provides extensive data on larval zebrafish locomotion, including at individual bout level with resolution of tail angles and bout frequency, and at a larger scale with distances, speeds, duration, and number of movements. Together, these assays establish a platform to evaluate the effect of photoswitchable and non-photoswitchable drugs in this animal model. The in-depth characterization of Carbadiazocine effects and its light-dependent activity posits it as a unique tool to study neural circuits underlying specific movements. These findings deepen our understanding of sensorimotor transformations and lay the foundations to probe native neuronal circuits in diverse animal species using patterns of photostimulation. Since neural computations are sensitive to perturbation in a highly spatiotemporally specific manner, tight control of circuit manipulation will be crucial for cracking the intricate networks underlying behavior.

## REFERENCES

[1] W. A. Velema, W. Szymanski, B. L. Feringa, “Photopharmacology: Beyond proof of principle” J. Am. Chem. Soc. 2014, 136, 2178–2191.

[2] K. Hu, J. Morstein, D. Trauner, “In Vivo Photopharmacology” 2018, DOI 10.1021/acs.chemrev.8b00037.

[3] J. Broichhagen, J. A. Frank, D. Trauner, “A roadmap to success in photopharmacology” Acc. Chem. Res. 2015, 48, 1947–1960.

[4] P. Kobauri, F. J. Dekker, W. Szymanski, B. L. Feringa, “Rational Design in Photopharmacology with Molecular Photoswitches” Angewandte Chemie 2023, e202300681.

[5] J. A. Frank, M.-J. Antonini, P.-H. Chiang, A. Canales, D. B. Konrad, I. C. Garwood, G. Rajic, F. Koehler, Y. Fink, P. Anikeeva, “In vivo photopharmacology enabled by multifunctional fibers” ACS Chem. Neurosci. 2020, 11, 3802–3813.

[6] V. A. Gutzeit, A. Acosta-Ruiz, H. Munguba, S. Häfner, A. Landra-Willm, B. Mathes, J. Mony, D. Yarotski, K. Börjesson, C. Liston, “A fine-tuned azobenzene for enhanced photopharmacology in vivo” Cell Chem. Biol. 2021, 28, 1648–1663.

[7] K. Hull, J. Morstein, D. Trauner, “In vivo photopharmacology” Chem. Rev. 2018, 118, 10710–10747.

[8] A. M. J. Gomila, P. Gorostiza, “In vivo applications of photoswitchable bioactive compounds” Molecular Photoswitches: Chemistry, Properties, and Applications, 2 Volume Set 2022, 811–842.

[9] G. Pietro da Silva, J. Doorduin, W. Szymanski, R. S. da Silva, P. Elsinga, C. D. Bonan, “Harnessing zebrafish as a model for photopharmacology: Insights into light-controlled biological effects of photoswitchable drugs” Drug Discov. Today 2025, 104477.

[10] C. A. MacRae, R. T. Peterson, “Zebrafish as tools for drug discovery” Nat. Rev. Drug Discov. 2015, 14, 721–731.

[11] C. Matera, A. M. J. Gomila, N. Camarero, M. Libergoli, C. Soler, P. Gorostiza, “Photoswitchable Antimetabolite for Targeted Photoactivated Chemotherapy” J. Am. Chem. Soc. 2018, 140, 15764–15773.

[12] D. Prischich, A. M. J. Gomila, S. Milla-navarro, G. Sangüesa, R. Diez-alarcia, B. Preda, C. Matera, L. Ramírez, E. Giralt, J. Hernando, J. J. Meana, P. De Villa, P. Gorostiza, D. Prischich, A. M. J. Gomila, S. Milla-navarro, G. Sangüesa, “Adrenergic Modulation with Photochromic Ligands” 2020, DOI 10.26434/chemrxiv.12203066.v1.

[13] C. Matera, P. Calvé, V. Casadó-Anguera, R. Sortino, A. M. J. Gomila, E. Moreno, T. Gener, C. Delgado-Sallent, P. Nebot, D. Costazza, “Reversible photocontrol of dopaminergic transmission in wild-type animals” Int. J. Mol. Sci. 2022, 23, 10114.

[14] R. Sortino, M. Cunquero, G. Castro-Olvera, R. Gelabert, M. Moreno, F. Riefolo, C. Matera, N. Fernàndez-Castillo, L. Agnetta, M. Decker, “Three-Photon Infrared Stimulation of Endogenous Neuroreceptors in Vivo” Angewandte Chemie International Edition 2023, 62, e202311181.

[15] L. Camerin, G. Maleeva, A. M. J. Gomila, I. Suárez-Pereira, C. Matera, D. Prischich, E. Opar, F. Riefolo, E. Berrocoso, P. Gorostiza, “Photoswitchable Carbamazepine Analogs for Non-Invasive Neuroinhibition In Vivo” Angewandte Chemie 2024, e202403636.

[16] D. Prischich, R. Sortino, A. Gomila-Juaneda, C. Matera, S. Guardiola, D. Nepomuceno, M. Varese, P. Bonaventure, L. de Lecea, E. Giralt, “In vivo photocontrol of orexin receptors with a nanomolar light-regulated analogue of orexin-B” Cellular and Molecular Life Sciences 2024, 81, 288.

[17] W. B. Barbazuk, I. Korf, C. Kadavi, J. Heyen, S. Tate, E. Wun, J. A. Bedell, J. D. McPherson, S. L. Johnson, “The syntenic relationship of the zebrafish and human genomes” Genome Res. 2000, 10, 1351–1358.

[18] K. Howe, M. D. Clark, C. F. Torroja, J. Torrance, C. Berthelot, M. Muffato, J. E. Collins, S. Humphray, K. McLaren, L. Matthews, “The zebrafish reference genome sequence and its relationship to the human genome” Nature 2013, 496, 498–503.

[19] C. B. Kimmel, W. W. Ballard, S. R. Kimmel, B. Ullmann, T. F. Schilling, “Stages of embryonic development of the zebrafish.” Dev. Dyn. 1995, 203, 253–310.

[20] C. B. Kimmel, S. L. Powell, W. K. Metcalfe, “Brain neurons which project to the spinal cord in young larvae of the zebrafish” Journal of Comparative Neurology 1982, 205, 112–127.

[21] E. A. Ober, H. A. Field, D. Y. R. Stainier, “From endoderm formation to liver and pancreas development in zebrafish” Mech. Dev. 2003, 120, 5–18.

[22] H. A. Field, E. A. Ober, T. Roeser, D. Y. R. Stainier, “Formation of the digestive system in zebrafish. I. Liver morphogenesis” Dev. Biol. 2003, 253, 279–290.

[23] G. N. Robertson, C. A. S. McGee, T. C. Dumbarton, R. P. Croll, F. M. Smith, “Development of the swimbladder and its innervation in the zebrafish, Danio rerio” J. Morphol. 2007, 268, 967–985.

[24] A. M. Stewart, O. Braubach, J. Spitsbergen, R. Gerlai, A. V Kalueff, “Zebrafish models for translational neuroscience research: from tank to bedside” Trends Neurosci. 2014, 37, 264–278.

[25] H. Feitsma, E. Cuppen, “Zebrafish as a cancer model” Molecular Cancer Research 2008, 6, 685–694.

[26] Y.-Y. Huang, “The optokinetic response in zebrafish and its applications” Frontiers in Bioscience 2008, 13, 1899.

[27] A. H. Burton, Q. Bai, E. A. Burton, “Sinusoidal analysis reveals a non-linear and dopamine-dependent relationship between ambient illumination and motor activity in larval zebrafish” Neurosci. Lett. 2021, 761, 136121.

[28] X. Gómez-Santacana, S. Pittolo, X. Rovira, M. Lopez, C. Zussy, J. A. R. Dalton, A. Faucherre, C. Jopling, J.-P. Pin, F. Ciruela, “Illuminating phenylazopyridines to photoswitch metabotropic glutamate receptors: from the flask to the animals” ACS Cent. Sci. 2017, 3, 81–91.

[29] D. Prischich, A. M. J. Gomila, S. Milla-Navarro, G. Sangüesa, R. Diez-Alarcia, B. Preda, C. Matera, M. Batlle, L. Ramírez, E. Giralt, “Adrenergic modulation with photochromic ligands” Angewandte Chemie International Edition 2021, 60, 3625–3631.

[30] A. V Kalueff, A. M. Stewart, R. Gerlai, “Zebrafish as an emerging model for studying complex brain disorders” Trends Pharmacol. Sci. 2014, 35, 63–75.

[31] D. A. Guggiana-Nilo, F. Engert, “Properties of the visible light phototaxis and UV avoidance behaviors in the larval zebrafish” Front. Behav. Neurosci. 2016, 10, 160.

[32] A. Bahl, F. Engert, “Neural circuits for evidence accumulation and decision making in larval zebrafish” Nat. Neurosci. 2020, 23, 94–102.

[33] R. Portugues, F. Engert, “The neural basis of visual behaviors in the larval zebrafish” Curr. Opin. Neurobiol. 2009, 19, 644–647.

[34] E. A. Naumann, J. E. Fitzgerald, T. W. Dunn, J. Rihel, H. Sompolinsky, F. Engert, “From whole-brain data to functional circuit models: the zebrafish optomotor response” Cell 2016, 167, 947–960.

[35] R. Tanaka, R. Portugues, “Algorithmic dissection of optic flow memory in larval zebrafish” Current Biology 2025, 35, 4870–4881.

[36] K. E. Fouke, Z. He, M. D. Loring, E. A. Naumann, “Neural circuits underlying divergent visuomotor strategies of zebrafish and Danionella cerebrum” Current Biology 2025, 35, 2457–2466.

[37] P. Simões, J. Moya-Díaz, L. Lagnado, “Quantifying the link between retinal performance and the optomotor response” Current Biology 2025, 35, 3908–3919.

[38] K. Bonnen, “Motion vision: Fish swimming to see” Current Biology 2023, 33, R30–R32.

[39] S. A. Budick, D. M. O’Malley, “Locomotor repertoire of the larval zebrafish: swimming, turning and prey capture” Journal of Experimental Biology 2000, 203, 2565–2579.

[40] K. A. Zalocusky, L. E. Fenno, K. Deisseroth, “Current challenges in optogenetics” Optogenetics 2013, 23–33.

[41] M. B. Orger, M. C. Smear, S. M. Anstis, H. Baier, “Perception of Fourier and non-Fourier motion by larval zebrafish” Nat. Neurosci. 2000, 3, 1128–1133.

[42] S. C. F. Neuhauss, O. Biehlmaier, M. W. Seeliger, T. Das, K. Kohler, W. A. Harris, H. Baier, “Genetic disorders of vision revealed by a behavioral screen of 400 essential loci in zebrafish” Journal of Neuroscience 1999, 19, 8603–8615.

[43] H. A. Burgess, M. Granato, “Modulation of locomotor activity in larval zebrafish during light adaptation” Journal of Experimental Biology 2007, 210, 2526–2539.

[44] R. E. Johnson, S. Linderman, T. Panier, C. L. Wee, E. Song, K. J. Herrera, A. Miller, F. Engert, “Probabilistic models of larval zebrafish behavior reveal structure on many scales” Current Biology 2020, 30, 70–82.

[45] E. I. Dragomir, V. Štih, R. Portugues, “Evidence accumulation during a sensorimotor decision task revealed by whole-brain imaging” Nat. Neurosci. 2020, 23, 85–93.

[46] H. Maaswinkel, L. Li, “Spatio-temporal frequency characteristics of the optomotor response in zebrafish” Vision Res. 2003, 43, 21–30.

[47] V. C. Fleisch, S. C. F. Neuhauss, “Visual behavior in zebrafish” Zebrafish 2006, 3, 191–201.

[48] D. L. Barabási, G. F. P. Schuhknecht, F. Engert, “Functional neuronal circuits emerge in the absence of developmental activity” Nat. Commun. 2024, 15, 364.

[49] M. Izquierdo-Serra, M. Gascón-Moya, J. J. Hirtz, S. Pittolo, K. E. Poskanzer, E. Ferrer, R. Alibés, F. Busqué, R. Yuste, J. Hernando, “Two-photon neuronal and astrocytic stimulation with azobenzene-based photoswitches” J. Am. Chem. Soc. 2014, 136, 8693–8701.

[50] S. Pittolo, H. Lee, A. Lladó, S. Tosi, M. Bosch, L. Bardia, X. Gómez-Santacana, A. Llebaria, E. Soriano, J. Colombelli, K. E. Poskanzer, G. Perea, P. Gorostiza, “Reversible silencing of endogenous receptors in intact brain tissue using 2-photon pharmacology” Proc. Natl. Acad. Sci. U. S. A. 2019, 116, 13680–13689.

[51] A. Garrido-Charles, M. Bosch, H. Lee, X. Rovira, S. Pittolo, A. Llobet, H. H.-W. Wong, A. Trapero, C. Matera, C. Papotto, “Photoswitching endogenous glutamate receptors in neural ensembles and single synapses in vivo” Brain Stimul. 2025.

