## supplemental information file for "Control of wildtype zebrafish optomotor response with a photoswitchable drug"

### Equivalent contribution

% Present address: Max Planck Institute for Brain Research, Frankfurt am Main, Germany

¥ Present address: Howard Hughes Medical Institute (HHMI), Cambridge, Massachusetts, United States

& Present address: Leibniz Institute for Neurobiology, Magdeburg, Germany

Keywords: Optomotor response (OMR), Circuit perturbation, Sensorimotor behavior, Photopharmacology, Photostimulation, Neuromodulation, Optogenetics, $Na_V$ voltage-gated sodium channels.

### 1 Materials and methods

#### 1.1 Zebrafish housing

Head-fixed experiments:

The larvae were obtained from crossing AB wild type zebrafish (*Danio Rerio*). Larvae were raised in groups of 20-30 individuals in filtered fish facility water in Petri dishes (9 cm diameter) daily cleaned and refilled. The animal development was checked every 24 hours, and abnormal or unhealthy embryos/larvae were removed. The environmental conditions during the development were the following: constant temperature at 28°C and 14h light – 10 h dark cycles. Larvae were fed daily with paramecia from at 4-day post fertilization (dpf). Experiments were performed with larvae at age range of 5-7 dpf. Sex cannot be determined at this stage. At the end of the experiments, larvae were euthanized in tricaine methanesulfonate 0.02%. All experiments were conducted with the approval of the Harvard University Standing Committee on the Use of Animals in Research and Training.

Free swimming experiments:

Wild-type zebrafish embryos (Tupfel long-fin strain, *Danio Rerio*) were obtained from the animal facility at the Barcelona Biomedical Research Park (PRBB). They were raised in darkness at 28 °C for 7 days in UV-filtered tap water, housed in Petri dishes that were cleaned and refilled daily. Embryonic development was monitored every 24 hours, and any unhealthy or abnormal embryos or larvae were removed and euthanized using 0.02% tricaine methanesulfonate. All experiments and procedures complied with the European Directive 2010/63/EU.

#### 1.2 Visual stimulus

Head-fixed experiments:

Random dot motion kinematograms (RDK) were employed as visual stimuli. Although various versions of this stimulus exist,[41] the one referenced in this work consists of a cloud of white dots (filtered, red) displayed against a black background. These stimuli was adopted from Bahl and Engert[26] and comprised approximately 1,000 dots, each 2 mm in diameter (generated online using custom-written programs in Python 3.7 and Panda3D), projected at 60 Hz using an AAXA P300 Pico Projector onto a circular arena with a 10 cm diameter, constructed from mildly light-scattering parchment paper. The white light from the projector was filtered with Hoya R62 (cut-on WL 620nm), 25.4mm Diameter, 2.5mm Thick, Colored Glass Longpass Filter to obtain red dots at wavelengths that are not interfering with the switching of Carbadiazocine. The behaviourally relevant visual field was approximately 5 cm in radius around the animal.

In the context of RDK, the term *coherence* refers to the strength of the stimulus, which can be regulated by varying the proportion of dots moving *coherently* in the same direction (Figure 3). A coherence level of 100% (the strongest stimulus) indicates that all dots move uniformly in one direction, while a coherence of 0% represents a static or flickering stimulus, where the dots flicker without directional movement. Coherence values between 0% and 100% reflect the probability that a dot follows the designated motion direction.

The random dots were projected in three different coherence levels (0, 25, 100%) each of them having two directions (-1, 1, left and right respectively). Before starting each trial, 5 seconds at 0% coherence were presented as a baseline stimulus (no movement, stationary flickering dots), then the stimulus is suddenly switching to one of the coherence levels, with the dots remaining stationary or translating continuously (speed: 1.8 cm $s^{-1}$) either leftwards or rightwards, perpendicular to the animal body. Closed loop was adopted, meaning that the visual stimulus is interrupted every time a bout is detected, ending the trail. A new trial can start only if the larvae are not responding during the initial 5 sec of adaptation, otherwise this baseline restarts. Each trial is lasting for a maximum of 45 sec and in absence of response from the fish (timeout) a new trial was automatically initiated. Stimulus protocol: 5 sec of 0% coherence, followed by 0%, 25% or 100% coherence moving rightward or leftward, followed by 5 s of 0% coherence. The stimulation order was random.

Only bouts within the threshold (bout start vigor: 50, end vigor: 10, custom-made tracking software, see section SI below) are detected while smaller swims are not registered and do not interrupt the trial. Each detected swim is labelled as either correct or incorrect based on the initial tail deflection or on the integration of the full bout. If the bout turns towards the same direction as the moving dots, it is classified as correct.

Each dot, whether static or moving, had a short average lifetime of 200 ms, during which it stochastically disappeared and immediately reappeared at a random location. This design ensured that the animals could not track individual dots, avoiding potential confounds in the experiments. The dots were red (> 620 nm, filtered from white light) against a black background (average scene luminance of approximately 120 Lux, Extech Instruments Light Meter LT300).

Free swimming experiments:

The stimuli consisted of 5 minutes in complete darkness followed by a pattern of blue and black stripes (10 in total) moving vertically at a speed of 10 revolutions per minute for 30 seconds in each direction (upwards and downwards) for 5 minutes. The stimuli was generated using OMR module in Zebrabox (Viewpoint Behavior Technology, France).

#### 1.3 Tracking and assay protocol

Head-fixed experiments:

Larvae were embedded in freshly made 2% agarose (UltraPure Low Melting Point Agarose, 16520-100, Invitrogen) at approximately 35°C, in non-treated Petri dishes (5 cm diameter). After solidifying (~ 1 min), the dish was filled with filtered facility water and the agarose around the tail and the mouth was removed leaving the minimum amount possible. The experiments were performed immediately after the embedding. The scene was illuminated from the bottom using infrared light-emitting diode (LED) panels (940 nm panel, Cop Security). Tracking was performed with a Grasshopper3-NIR camera (FLIR Systems) equipped with a Navitar Zoom 7000 lens (18–108 mm) and an R72 long-pass filter (Hoya). Using a custom-written Python 3.7-based software package, the trail was tracked following 27 nodes along the tail. Tracking was conducted using two computers, each connected to four cameras and two projectors, enabling independent stimulation and closed-loop tracking of eight individual fish simultaneously.

Larvae were divided into three groups: (1) fish treated with 10 µM or 30 µM solution of Carbadiazocine in its dark thermally relaxed isoform (100% *cis*), (2) fish treated with Carbadiazocine pre-illuminated solution at 405 nm (50% *trans*) and (3) vehicles (fish water with 0.1% and 0.3% DMSO respectively). Experiments started at least 10 minutes after embedding. For all three groups of head-fixed larvae, basal swimming activity and responses to visual stimuli were recorded for 1 hour in filtered facility water. This duration ensured the collection of a sufficient number of trials for each coherence level. Subsequently, the filtered facility water was removed from the Petri dish using a plastic pipette, and the respective solutions for each group were added. The larvae were incubated in the new solutions for 20 minutes before being exposed to the visual stimuli and tracked for one additional hour.

Free swimming experiments:

Free-swimming Optomotor Response (OMR) assay was performed and recorded using Zebrabox device and analyzed with Zebralab software (Viewpoint, France). In this case, the larvae were placed in 10-well plates containing 2 mL of fish water per well. The stimuli screen was below the zebrafish larvae plate inside the Zebrabox. The protocol of the assay consisted of 5 minutes in complete darkness followed by 5 cycles of stimuli as described above. Basal response was measured and afterwards larvae were left to rest for 20 minutes. Then, 1 mL was removed from the well to add 1mL of either vehicle (DMSO 0.1 %), or Carbadiazocine 10 µM in dark conditions or pre-illuminated for 10 minutes with a 400 nm LED (final concentrations in well were 0.05% DMSO or 5 µM Carbadiazocine in 0.05% DMSO). Immediately after the addition of the corresponding compound, the plate was placed in the Zebrabox to measure again the OMR response.

The recordings obtained from each experiment were processed through two different modules of the Zebralab software to obtain the XY coordinates of each larva and the number of movement events, swam distance and time spent swimming at different speeds, classified as slow (< 2 $mm \cdot s^{-1}$), intermediate (2-6 $mm \cdot s^{-1}$) and fast (> 6 $mm \cdot s^{-1}$). Data was integrated every 1 second and 30 seconds.

OMR assay correctness was analyzed by comparing the Delta Y coordinate for each larva during the 30 second bins of the stimuli period (10 in total, 5 with stripes moving upwards and 5 downwards). A correct response was considered when the larva Delta Y for the bin was in accordance with the direction of the stimuli (i.e., if the stripes were moving upwards and the final position of the larvae was upper than the initial) and above a Delta Y threshold established as the mean Delta Y of control larvae (basal condition) in the absence of stimuli in 30 seconds bins in basal condition. In this way, we ensure that when we count a positive response the direction of the movement was correct and not random but guided by the stripes. Other thresholds were evaluated to assess correctness, such as an average Delta Y for each plate in basal condition in the absence of stimuli, or individually for each larva, yielding the same results (see Figure S16).

#### 2 Additional analysis for head-fixed larval zebrafish

Data from the correctness in head-fixed experiments were normalized to their respective basal performance for each coherence level (0%, 25% and 100%) and treatment (vehicle 0.3% DMSO, dark Carbadiazocine 30 µM and 405 nm pre-illuminated Carbadiazocine 30 µM). A significant drop in the percentage of correctness was observed for coherence 100% for pre-illuminated Carbadiazocine compared to vehicle and dark Carbadiazocine.

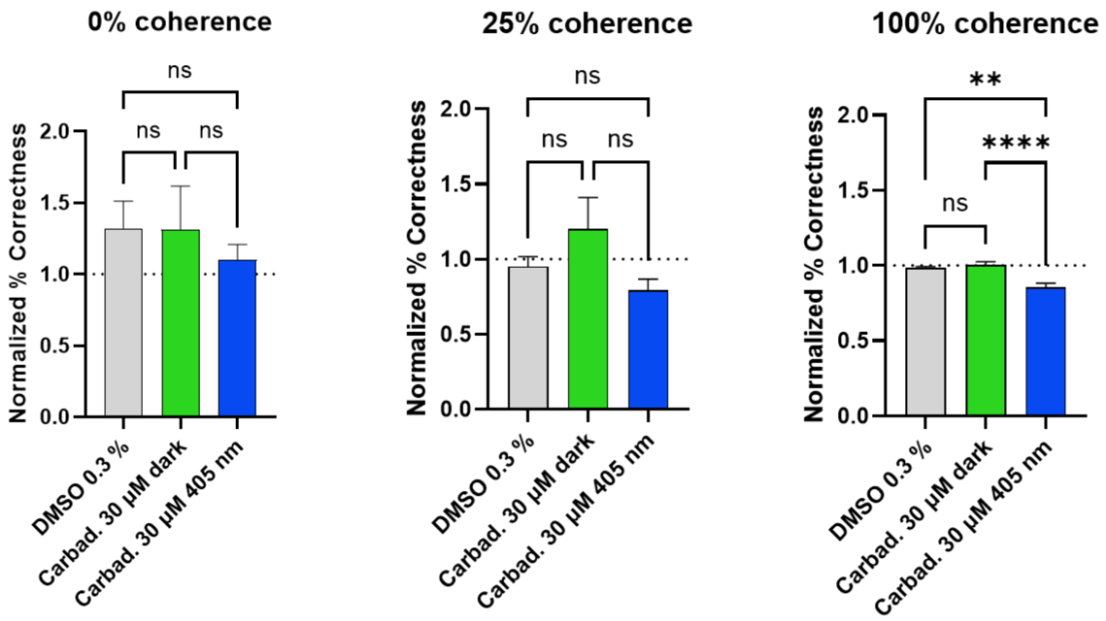

**Figure S1:** Normalized percentage of correctness. For each level of coherence (0%, 25% and 100%) and for each respective treatment (vehicle DMSO 0.3%, dark Carbadiazocine at 30 µM and 405 nm pre-illuminated Carbadiazocine at 30 µM) the data were normalized to the correctness at the basal performance in fish facility water. **** $p<0.0001$ and ** $p=0.0011$ One-Way ANOVA with Kruskal-Wallis multiple comparison test. Error bars represent S.E.M.

The experiment with Carbadiazocine at 10 µM only showed significance for the pre-illuminated isomer at 100% coherence, most likely due to the dilution of the solution below the effective concentration caused by the presence of agarose.

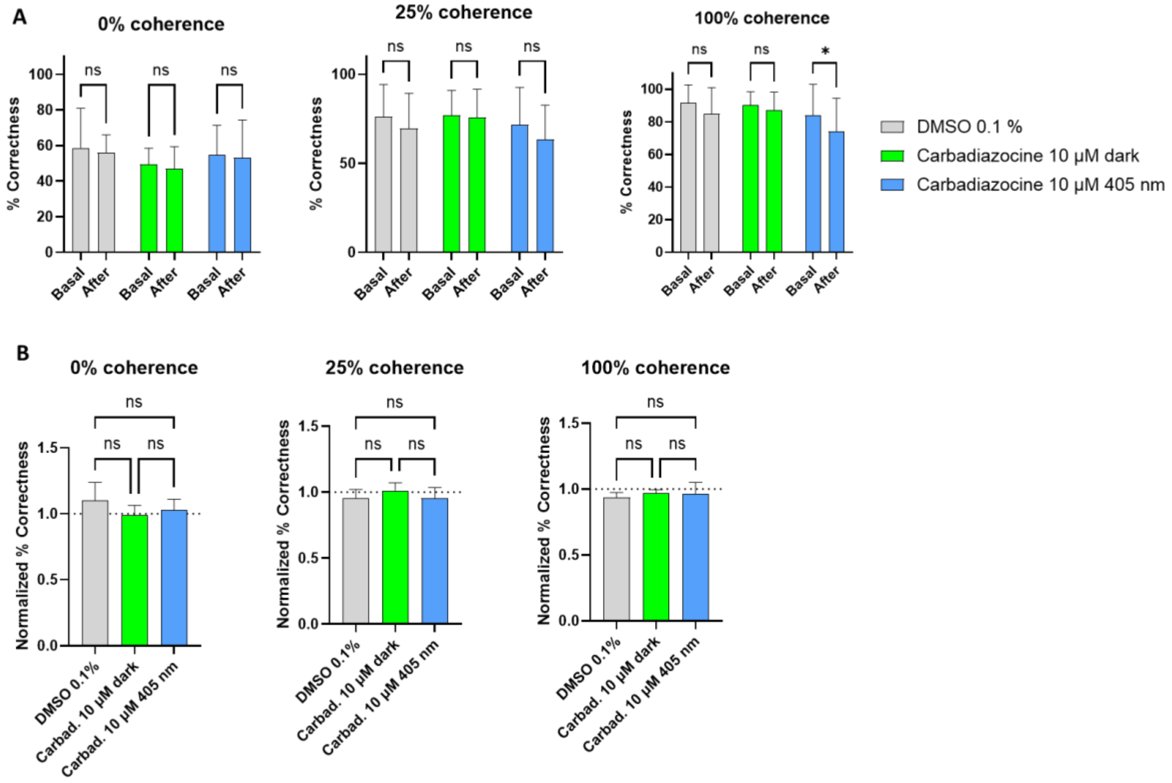

**Figure S2:** Results for correctness for zebrafish larvae treated with Carbadiazocine at 10 µM. **A)** Percentage of correctness for each level of coherence (0%, 25% and 100%). Significant drop in correctness was only observed for 100% coherence with larvae treated with a solution of 405 nm pre-illuminated Carbadiazocine. "Basal" represents the fish performance in fish facility water; "after" indicates the fish performance post addition of the prepared solutions (vehicle 0.1% DMSO, dark Carbadiazocine at 10 µM and 405 nm pre-illuminated Carbadiazocine at 10 µM). * $p<0.05$ Two-Way ANOVA with Šídák's multiple comparisons test. Error bars represent S.E.M. **B)** Normalized percentage of correctness. For each level of coherence (0%, 25% and 100%) and for each respective treatment (vehicle DMSO 0.1%, dark Carbadiazocine at 10 µM and 405 nm pre-illuminated Carbadiazocine at 10 µM) the data were normalized to the correctness at the basal performance in fish facility water. One-Way ANOVA with Kruskal-Wallis multiple comparison test, ns = not significant. Error bars represent S.E.M.

The following parameters were analysed and represented in the graphs below: time to bout, number and fraction of completed trials, percentage correct over time. We considered these results less representative compared to the correctness.

The interbout-intervals (time to bout) were shortened with 405 nm pre-illuminated Carbadiazocine compared to vehicle and dark compound. This behaviour can be observed from the recorded traces or the recorded video from the camera, but it is not fairly represented in "time to bout". In fact, quantifying this parameter cannot be done with accuracy considering that the tracking software only detected as bouts the tail movements above the manually pre-set threshold (see Methods). Larvae exposed to 405 nm pre-illuminated Carbadiazocine exhibited continuous small swims rather than distinct bouts. Given the design of the visual stimuli, counting these small swims as bouts would prevent new trials from starting, meaning the animals would only experience the initial 5 seconds of the 0 % coherence baseline. As a result, in the time-to-bout quantification, all data related to small swims are lost, and only detected bouts that surpass the threshold are recorded. For this reason, we do not consider the analysis of this parameter to be fully representative.

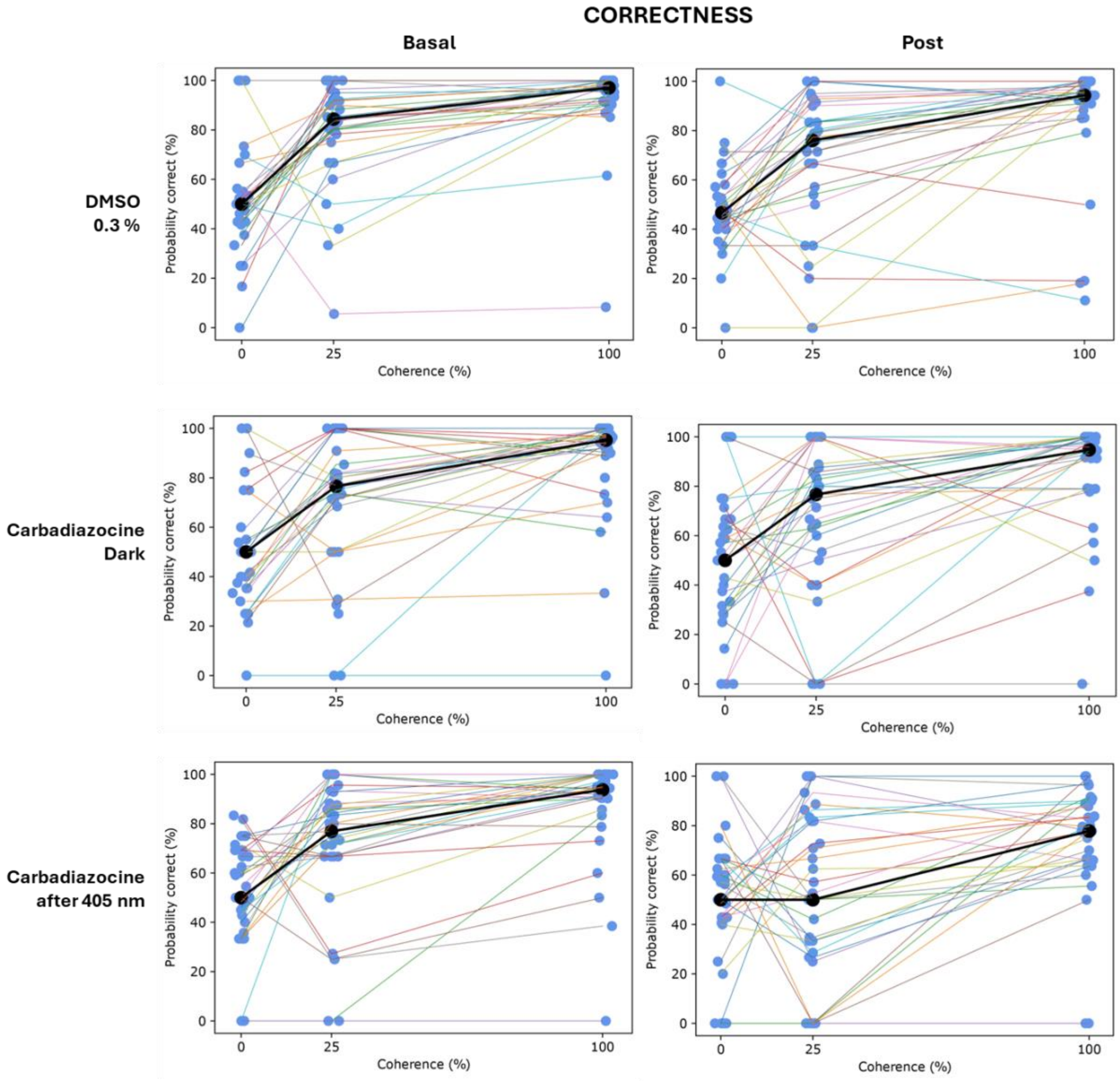

**Figure S3:** Percentage of correctness for the three levels of coherence (0-25-100%) and for treatments (vehicle 0.3% DMSO, dark-adapted Carbadiazocine at 30 µM and 405 nm pre-irradiated Carbadiazocine at 30 µM). "Basal" graphs correspond to the performance in filtered fish water, "post" graphs correspond to the performance after the addition of the respective treatment solutions. Blue dots represent the average % of correctness (in successful trials) for each larva. Colored lines are connecting the average performance of each larva among the different levels of coherence. Black dots show the average % of correctness among all the larvae, for each level of coherence.

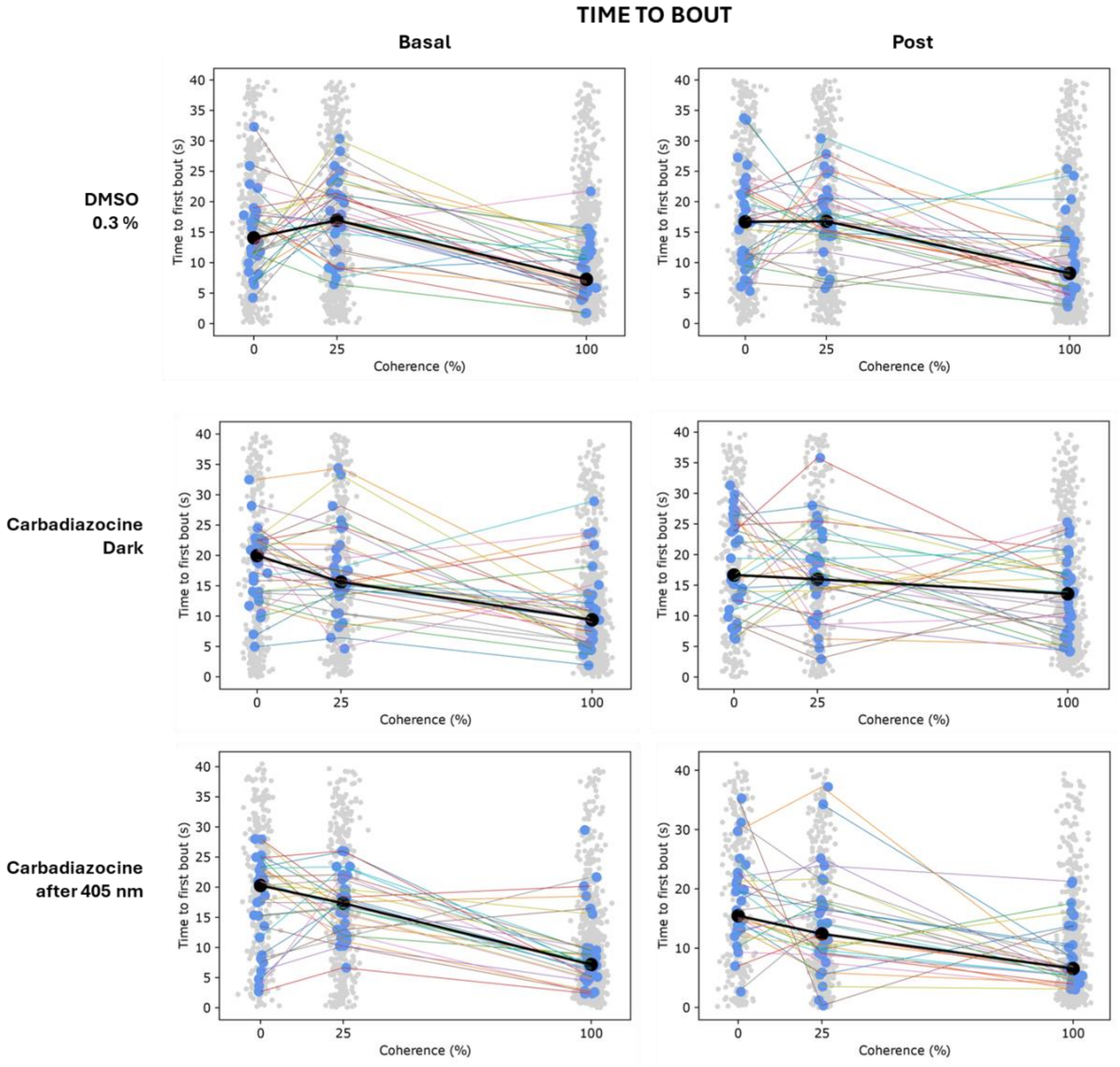

**Figure S4:** Time to first bout (s) for the three level of coherence (0-25-100%) and for the treatments (vehicle 0.3% DMSO, dark-adapted Carbadiazocine at 30 µM and 405 nm pre-irradiated Carbadiazocine at 30 µM). "Basal" graphs correspond to the performance in filtered fish water, "post" graphs correspond to the performance after the addition of the treatment solutions. Blue dots represent the average time to bout (in successful trials) for each larva. Grey dots represent each data point. Coloured lines are connecting the average performance of each larva among the different levels of coherence. Black dots show the average time to bout among all the larvae, for each level of coherence.

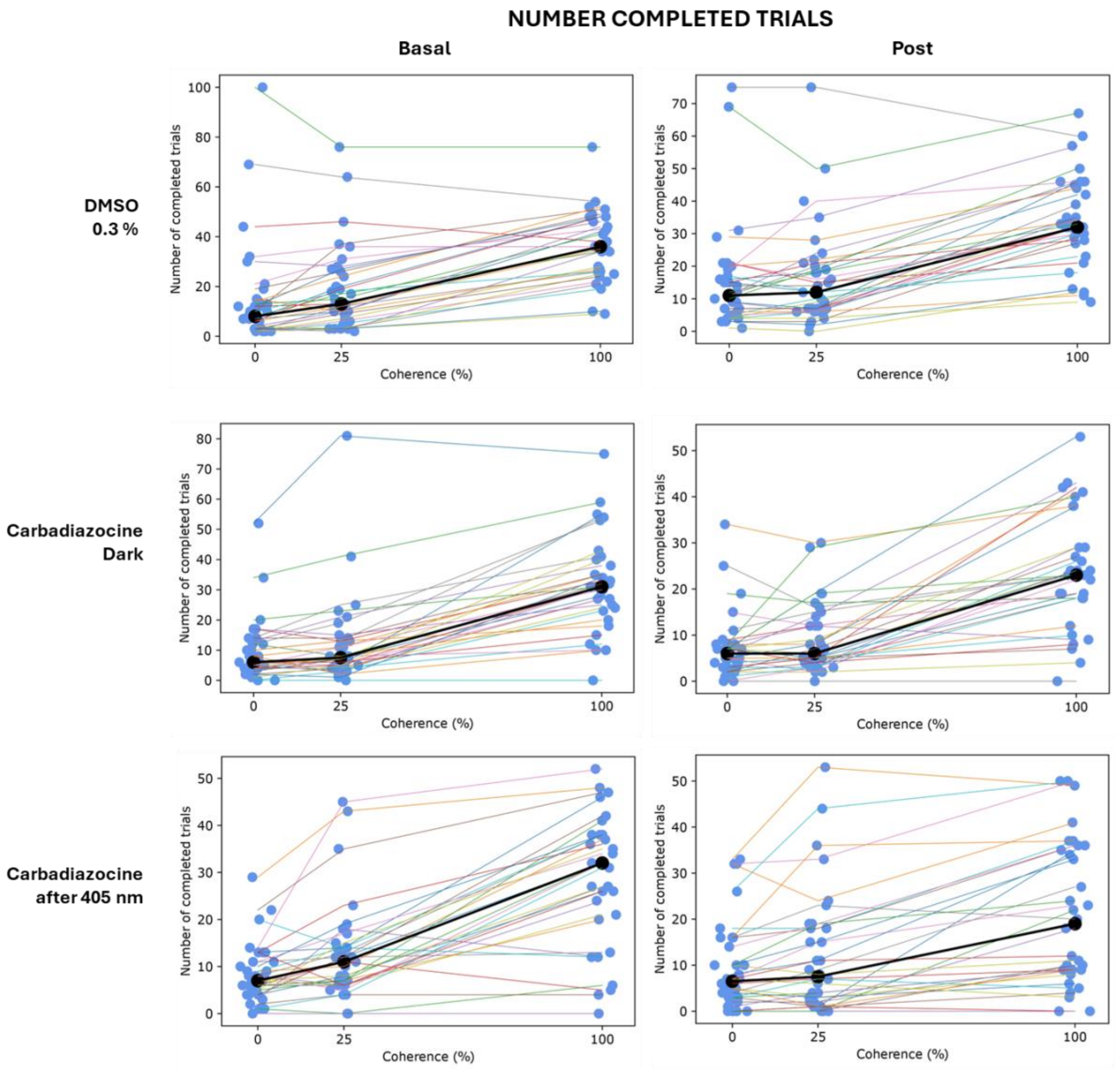

**Figure S5:** Number of completed trials for the three level of coherence (0-25-100%) and for the three treatments used (vehicle 0.3% DMSO, dark-adapted Carbadiazocine at 30 µM and 405 nm pre-irradiated Carbadiazocine at 30 µM). "Basal" graphs correspond to the performance in filtered fish water, "post" graphs correspond to the performance after the addition of the treatment solutions. Blue dots represent the average number of completed trials for each larva. Colored lines are connecting the average performance of each larva among the different levels of coherence. Black dots show the average number of completed trials among all the larvae, for each level of coherence.

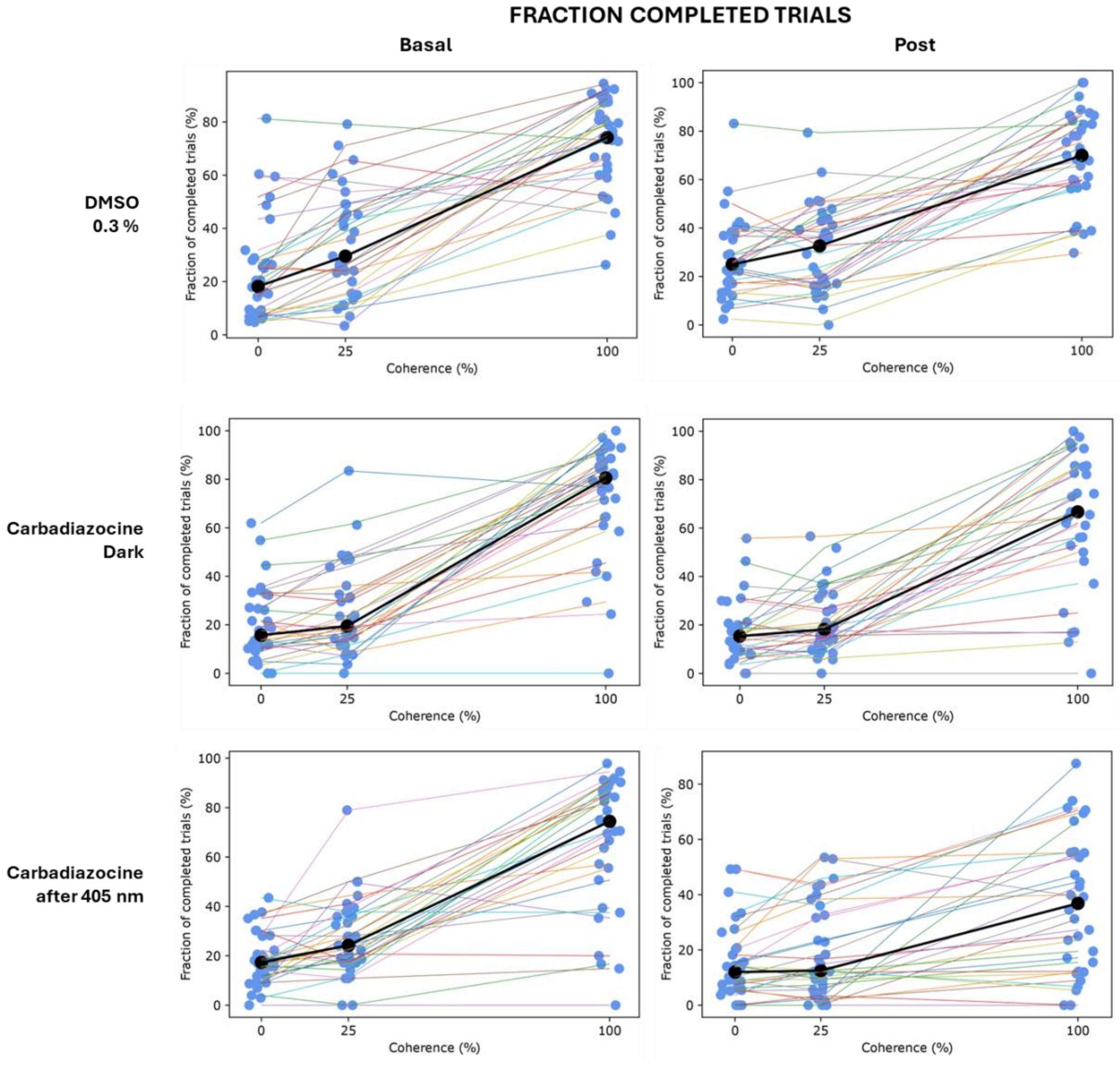

**Figure S6:** Fraction of completed trials (% completed trials/total trials) for the three level of coherence (0-25-100%) and for the three treatments used (vehicle 0.3% DMSO, dark-adapted Carbadiazocine at 30 µM and 405 nm pre-irradiated Carbadiazocine at 30 µM). "Basal" graphs correspond to the performance in filtered fish water, "post" graphs correspond to the performance after the addition of the treatment solutions. Blue dots represent the average fraction of completed trials for each larva. Colored lines are connecting the average performance of each larva among the different levels of coherence. Black dots show the average fraction of completed trials among all the larvae, for each level of coherence.

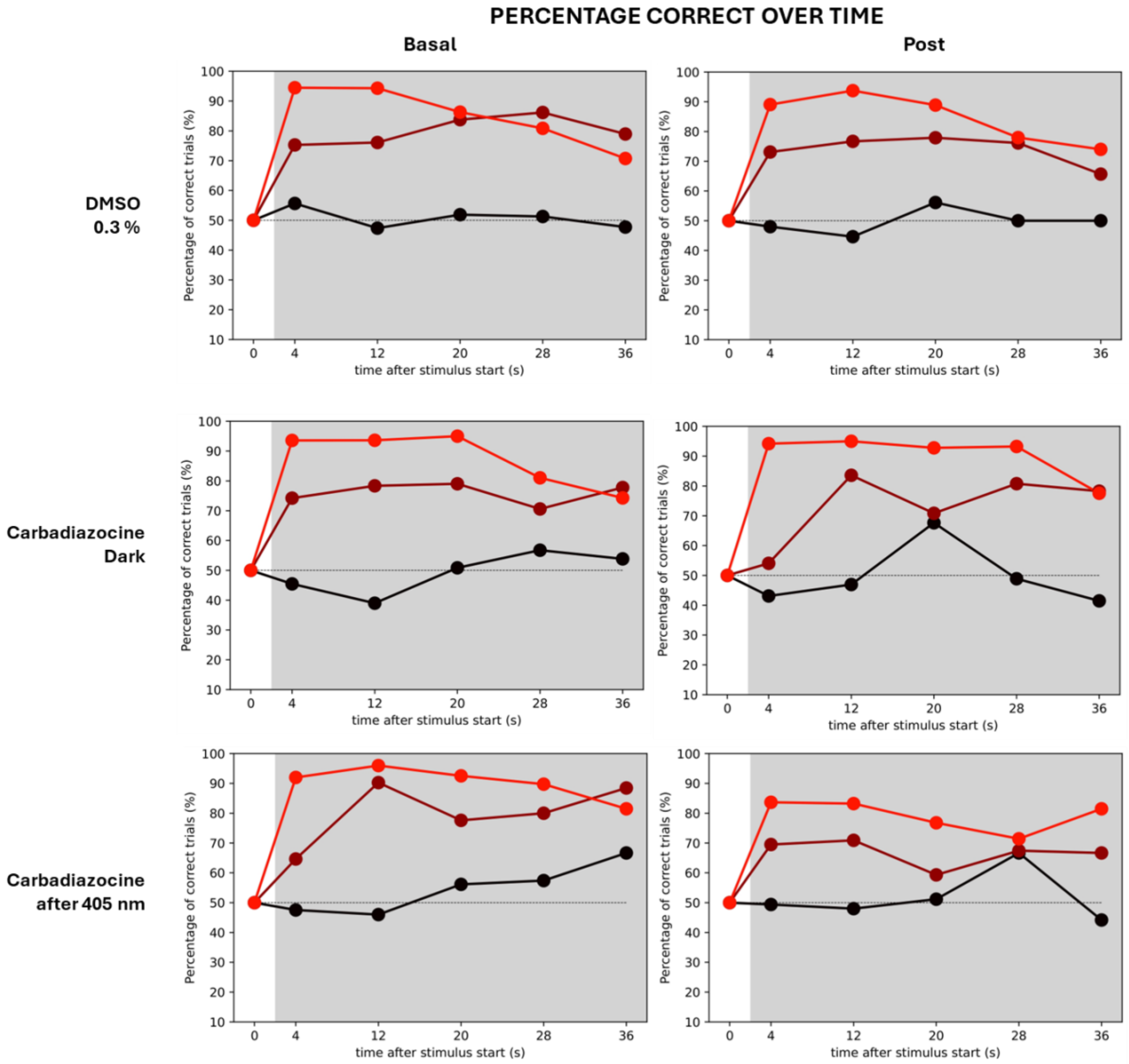

Figure S7: Percentage correct over time (%) for the three level of coherence (0-25-100%) and for the treatments used (vehicle 0.3% DMSO, dark-adapted Carbadiazocine at 30 µM and 405 nm pre-irradiated Carbadiazocine at 30 µM). "Basal" graphs correspond to the performance in filtered fish water, "post" graphs correspond to the performance after the addition of the treatment solutions. Light red line: 100% coherence; Dark red line: 25% coherence; Black line: 0% coherence. The length of the stimuli has been divided into bin times (0-4 sec, 4-12 sec, 12-20 sec, 20-28 sec and 28-36 sec). For each bin time it is shown the average % of correctness (dots) for the larvae responding to the stimuli during each bin time frame, for each level of coherence.

#### TRACES

**A** **DMSO 0.3 %**

**Basal**

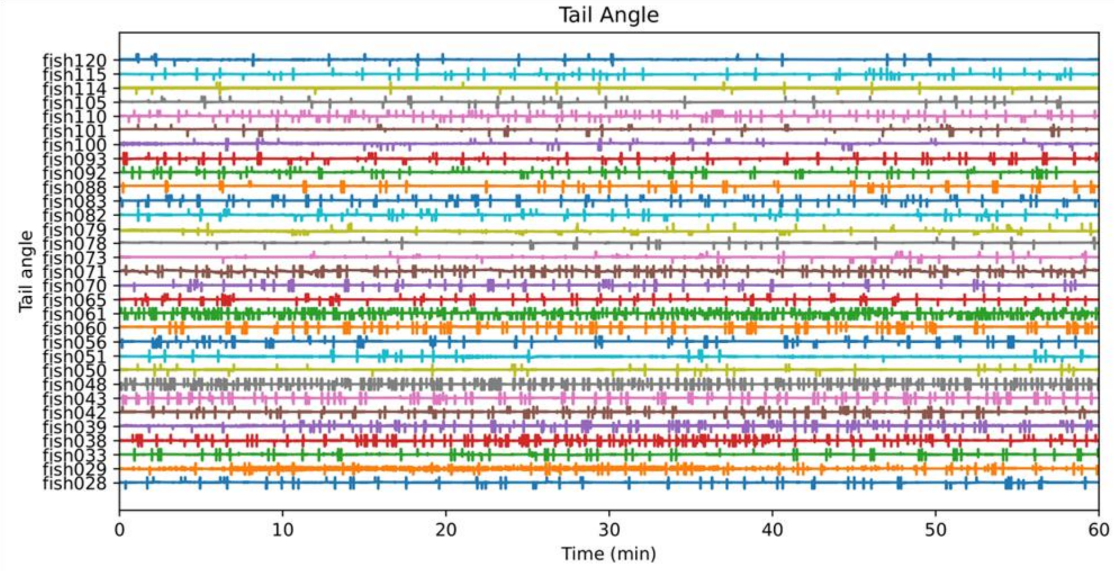

**Post**

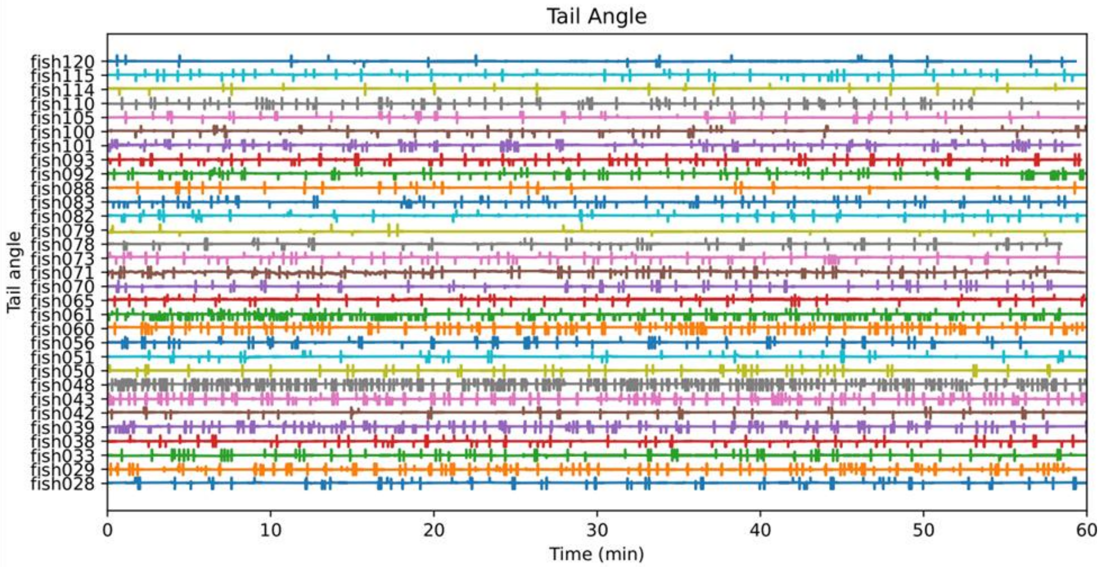

**B**

**Carbadiazocine Dark**

**Basal**

**Post**

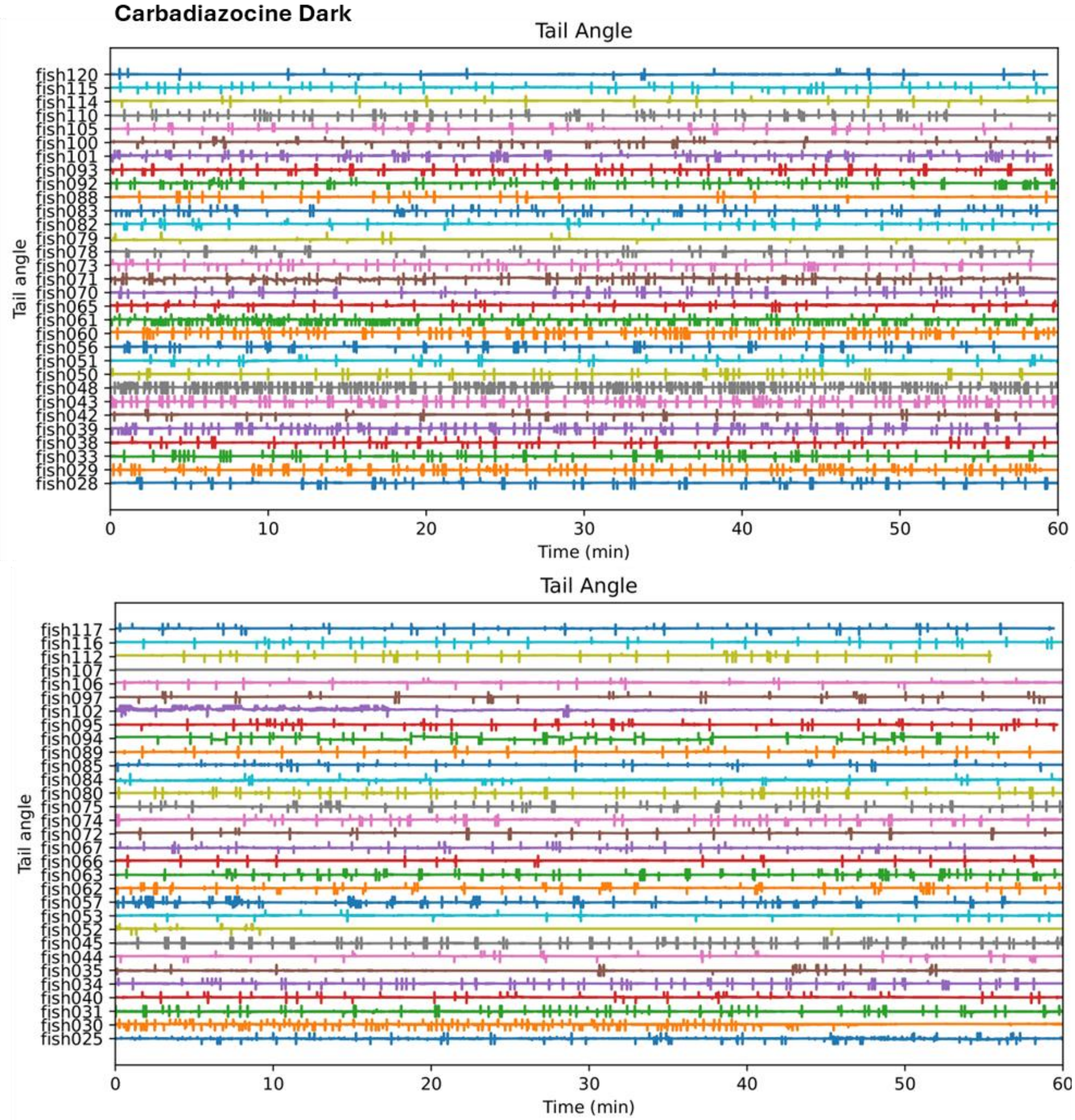

**C**

**Basal**

**Post**

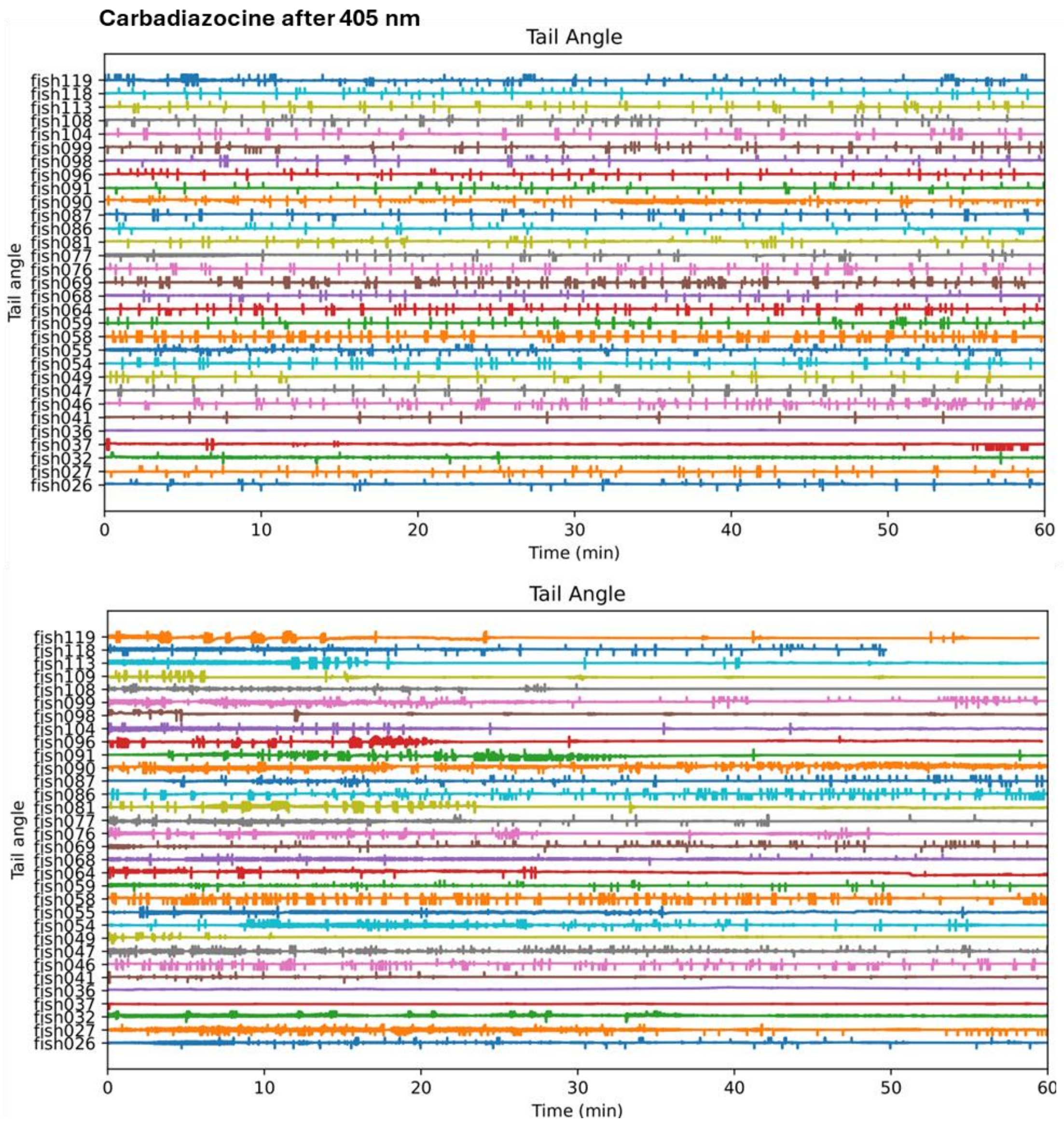

**Figure S8:** Raw traces recording the tracked swimming activity for each head-fixed larva. During the recordings the larvae were exposed to the visual stimuli, although it is not integrated into the row traces graphs. “Basal” graphs correspond to the performance in filtered fish water, “post” graphs correspond to the performance after the addition of the treatment solutions. Upper lines indicate “right” tail deflection, lower lines indicate “left” tail deflection **A)** Basal and post traces for vehicle (0.3% DMSO). **B)** Basal and post traces for the solution of dark-adapted Carbadiazocine isoform at 30 µM. **C)** Basal and post traces for the solution of 405 nm pre-illuminated Carbadiazocine at 30 µM.

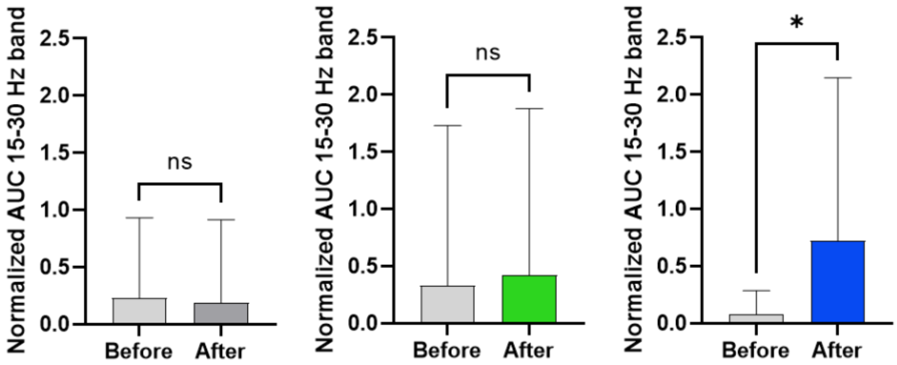

**Figure S9:** Normalized Area under the curve (AUC) of the power spectrum of subthreshold swims shown in Figure 3.C for head-fixed larvae treated with vehicle 0.3% DMSO (dark grey), dark-adapted Carbadiazocine at 30 µM (green) and 405 nm pre-irradiated Carbadiazocine at 30 µM (blue). "Before" bars correspond to the performance in filtered fish water, "After" bars correspond to the performance after the addition of the treatment solutions. AUC for the 15-30 Hz frequency band was normalized by the total AUC of the power spectrum. The contribution of this frequency movements is significantly increased upon treatment with pre-illuminated Carbadiazocine. Data is represented as mean ± standard deviation. *$p<0.05$; ns = not significant, Wilcoxon matched pairs signed rank test.

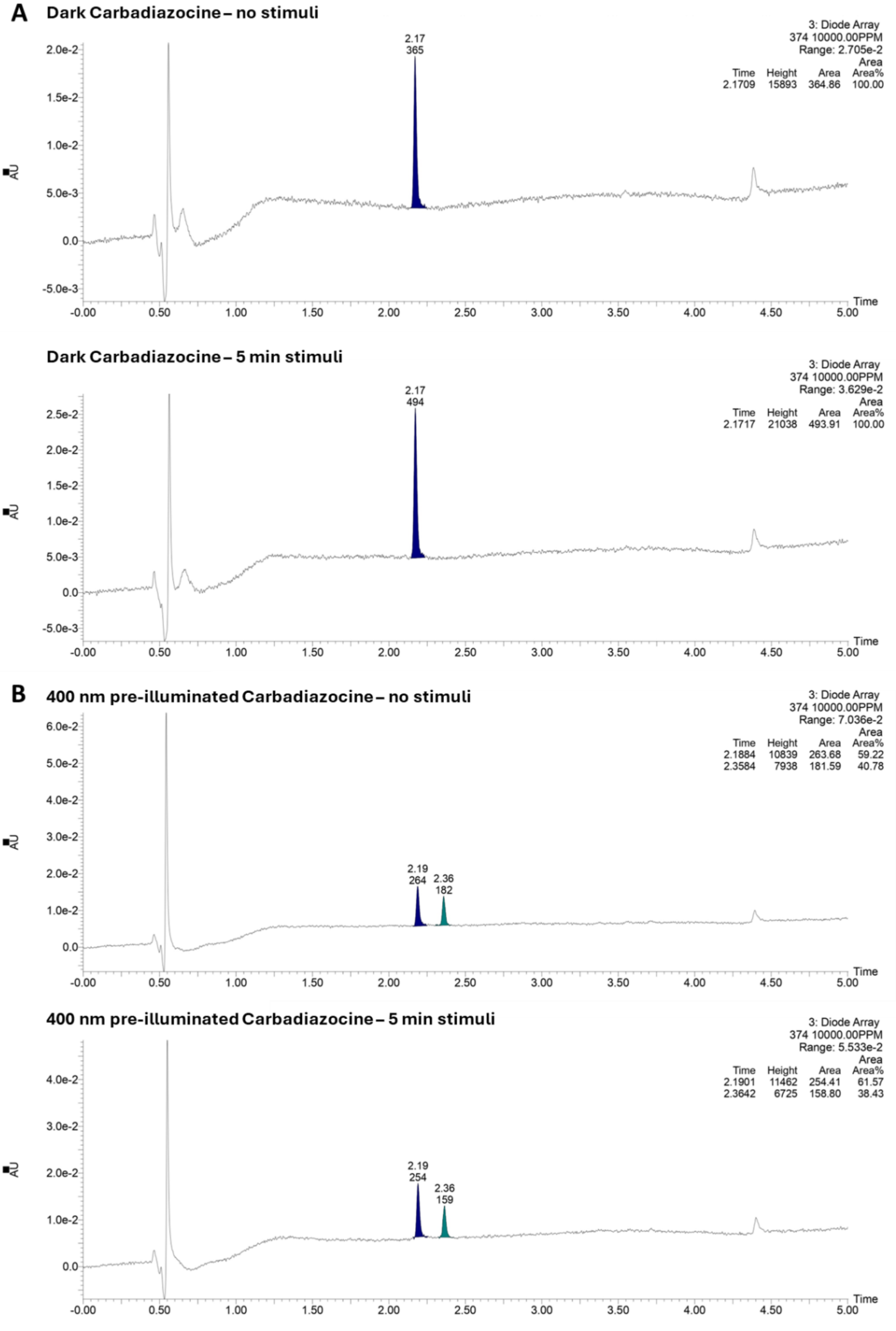

A
Dark Carbadiazocine – no stimuli
3: Diode Array
374 10000.00PPM
Range: 2.705e-2
Time Height Area Area%
2.1709 15893 364.86 100.00
2.17
365
Dark Carbadiazocine – 5 min stimuli
3: Diode Array
374 10000.00PPM
Range: 3.629e-2
Time Height Area Area%
2.1717 21038 493.91 100.00
2.17
494
B
400 nm pre-illuminated Carbadiazocine – no stimuli
3: Diode Array
374 10000.00PPM
Range: 7.036e-2
Time Height Area Area%
2.1884 10839 263.68 59.22
2.3584 7938 181.59 40.78
2.19
264
2.36
182
400 nm pre-illuminated Carbadiazocine – 5 min stimuli
3: Diode Array
374 10000.00PPM
Range: 5.533e-2
Time Height Area Area%
2.1901 11462 254.41 61.57
2.3642 6725 158.80 38.43
2.19
254
2.36
159
AU
Time

**Figure S10:** HPLC photostationary quantification. **A)** Dark Carbadiazocine at 5 µM remains 100% cis upon exposure to blue stripes visual stimuli. **B)** 400 nm pre-illumined Carbadiazocine at 5 µM does not significantly reduce its *trans* % isomer upon exposure to blue stripes visual stimuli (41% vs 38%).

### 3 Additional analysis for free-swimming larval zebrafish

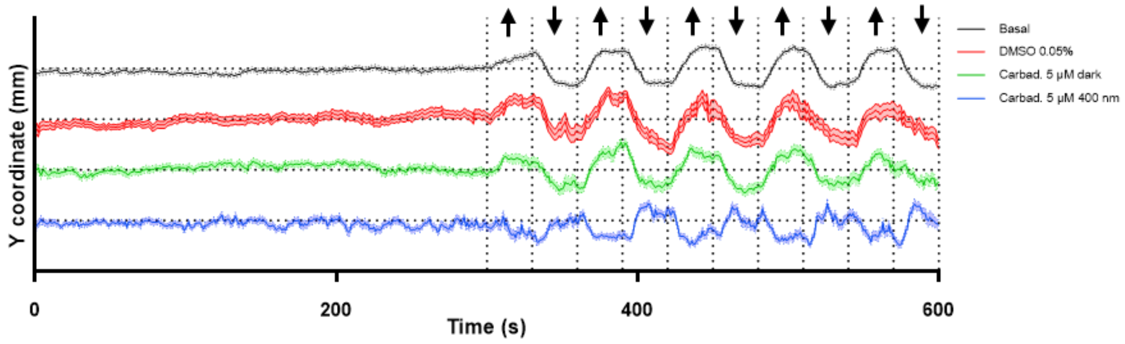

**Figure S11:** Optomotor response assay Y coordinate through time for larvae in basal condition, treated with vehicle (DMSO 0.05 %), dark Carbadiazocine 5 µM and 400 nm pre-illuminated Carbadiazocine 5 µM, here represented as stacked lines. Horizontal dotted lines represent the central position of the well, whereas vertical dotted lines represent the timepoint where the stimuli direction changes, indicated by the arrows. Treatment with 400 nm pre-illuminated Carbadiazocine affects the capability of following the stripes direction correctly.

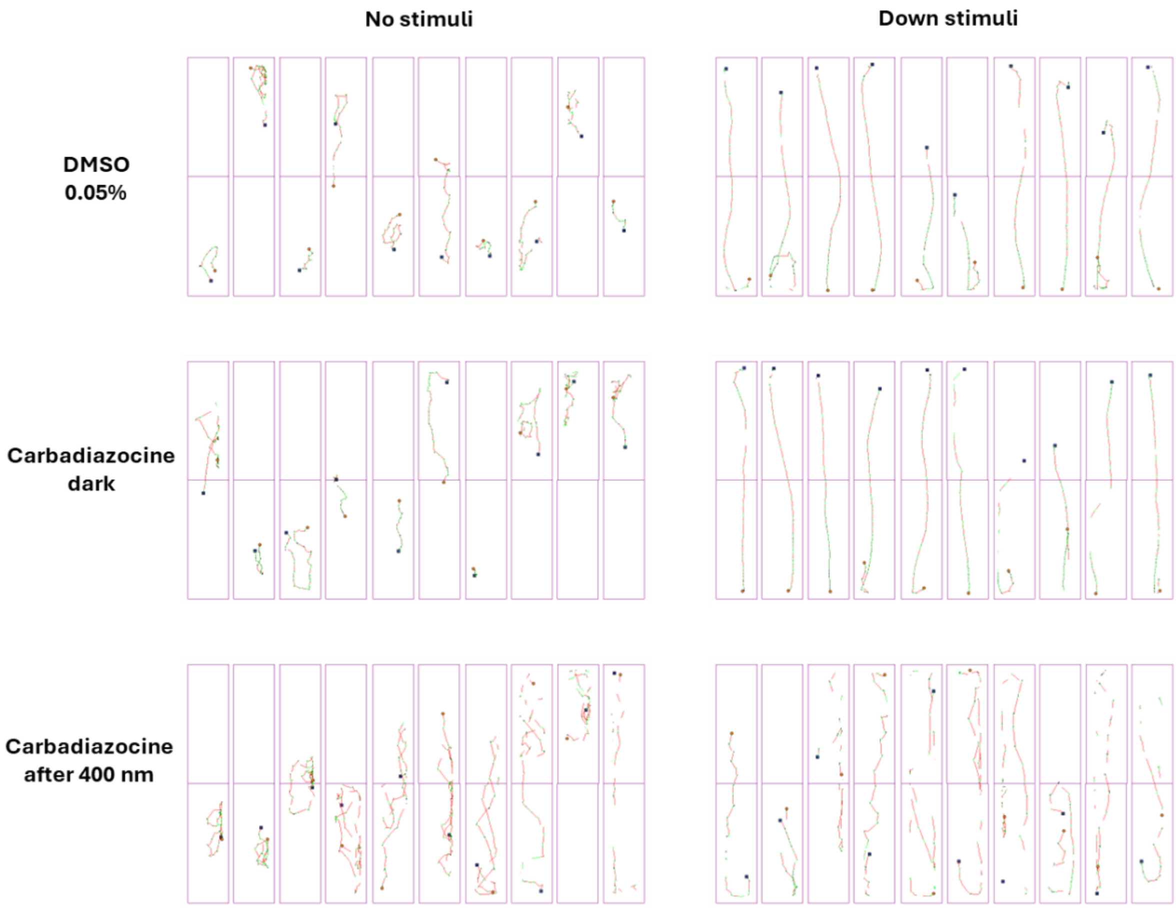

**Figure S12:** Examples of raw traces of swimming activity during a 30 sec time bin in free swimming larvae for each treatment condition (vehicle 0.05% DMSO, dark-adapted Carbadiazocine isoform at 5 µM, 400 nm pre-illuminated Carbadiazocine at 5 µM). In the left panel "No stimuli" corresponds to the swimming activity without visual stimulation. In the right panel "Down stimuli" corresponds to the swimming activity in presence of stripes moving downwards. Squares indicate the initial position of the larvae. Dots represent the final position of the larvae. Red lines correspond to fast movements (distance travelled at ≥ 6 $mm{\cdot}s^{-1}$); green lines correspond to distance travelled at 2–6 $mm{\cdot}s^{-1}$; black lines correspond to distance travelled at ≤ 2 $mm{\cdot}s^{-1}$.

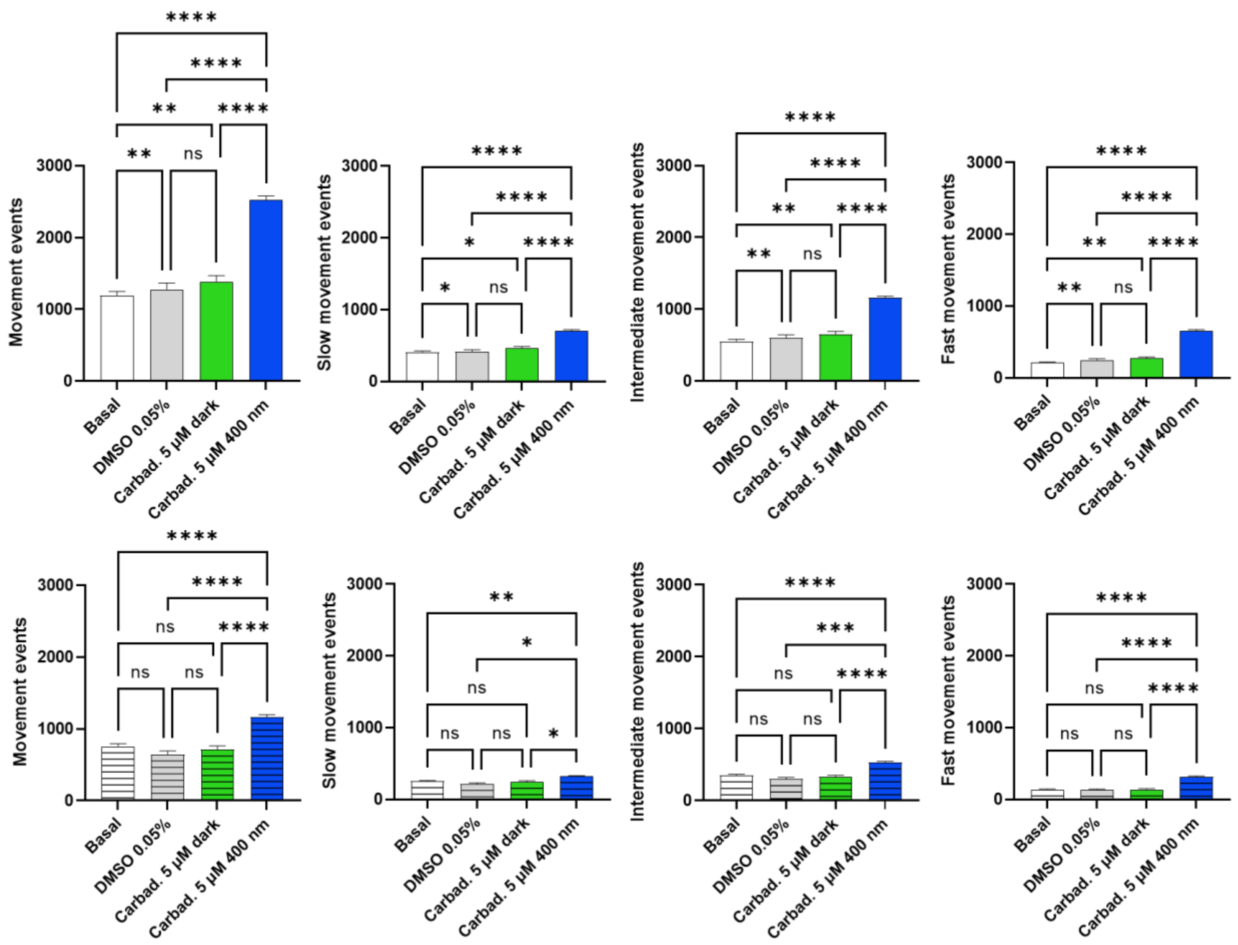

**Figure S13:** Count of total movement events (i.e. every time a larva initiates a movement) and classified by speed of movement for larvae in basal condition, treated with vehicle (DMSO 0.05 %), benchtop Carbadiazocine and 400 nm pre-illuminated Carbadiazocine. Smooth bars represent the count for the complete experiment and striped bars during the stimuli period. Data represented as mean ± S.E.M. ****p<0.0001; ***p<0.001; **p<0.01; *p<0.05; ns = not significant, Mixed-effect analysis followed by Tukey's multiple comparisons test.

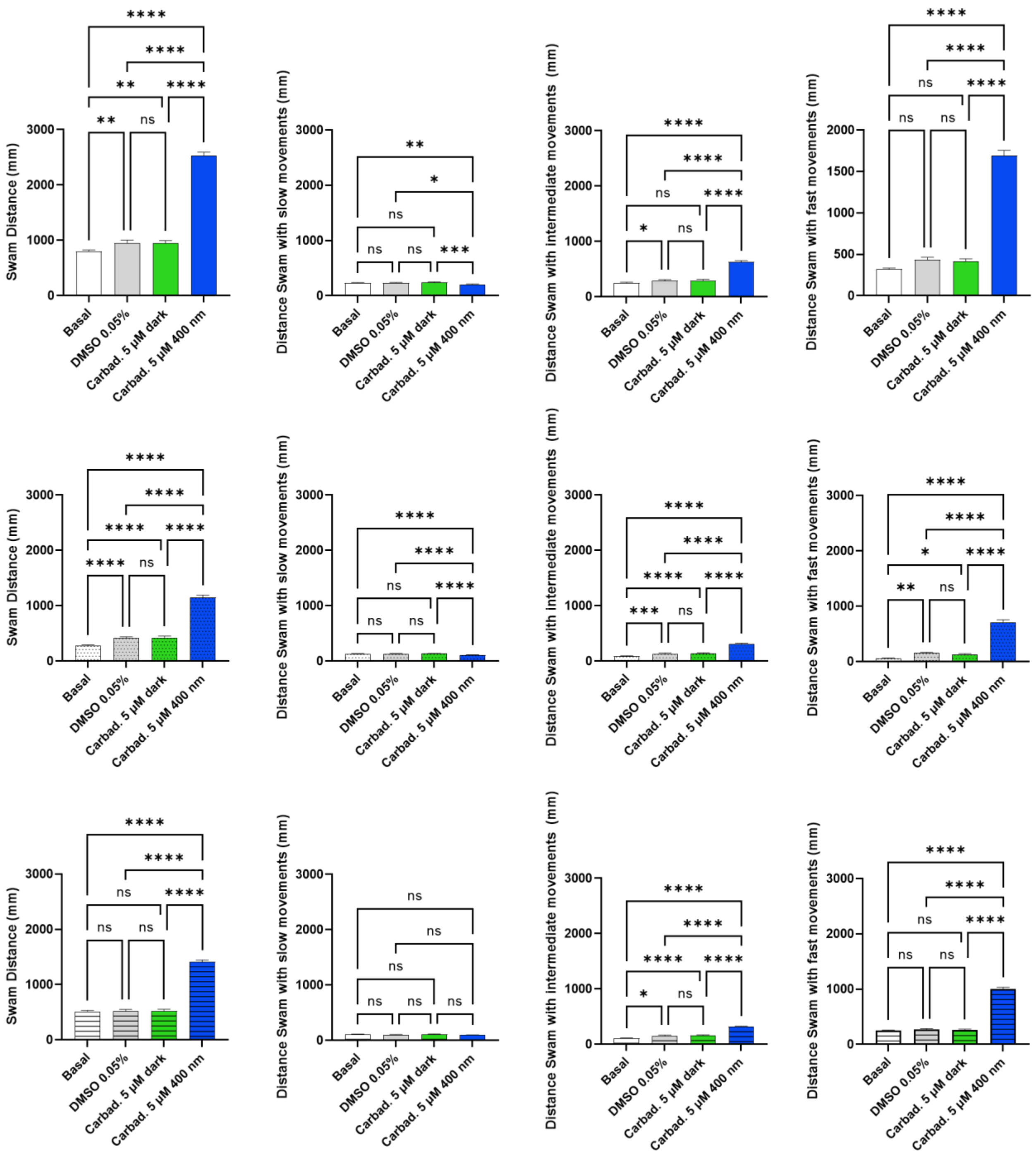

**Figure S14:** Total swam distances and classified by speed of movement for larvae in basal condition, treated with vehicle (DMSO 0.05 %), benchtop Carbadiazocine and 400 nm pre-illuminated Carbadiazocine. Smooth bars represent the swam distances for the complete experiment, dotted bars during the no-stimuli period and striped bars during the stimuli period. Data represented as mean ± S.E.M. ****p<0.0001; **p<0.01; *p<0.05; ns = not significant, Mixed-effect analysis followed by Tukey's multiple comparisons test.

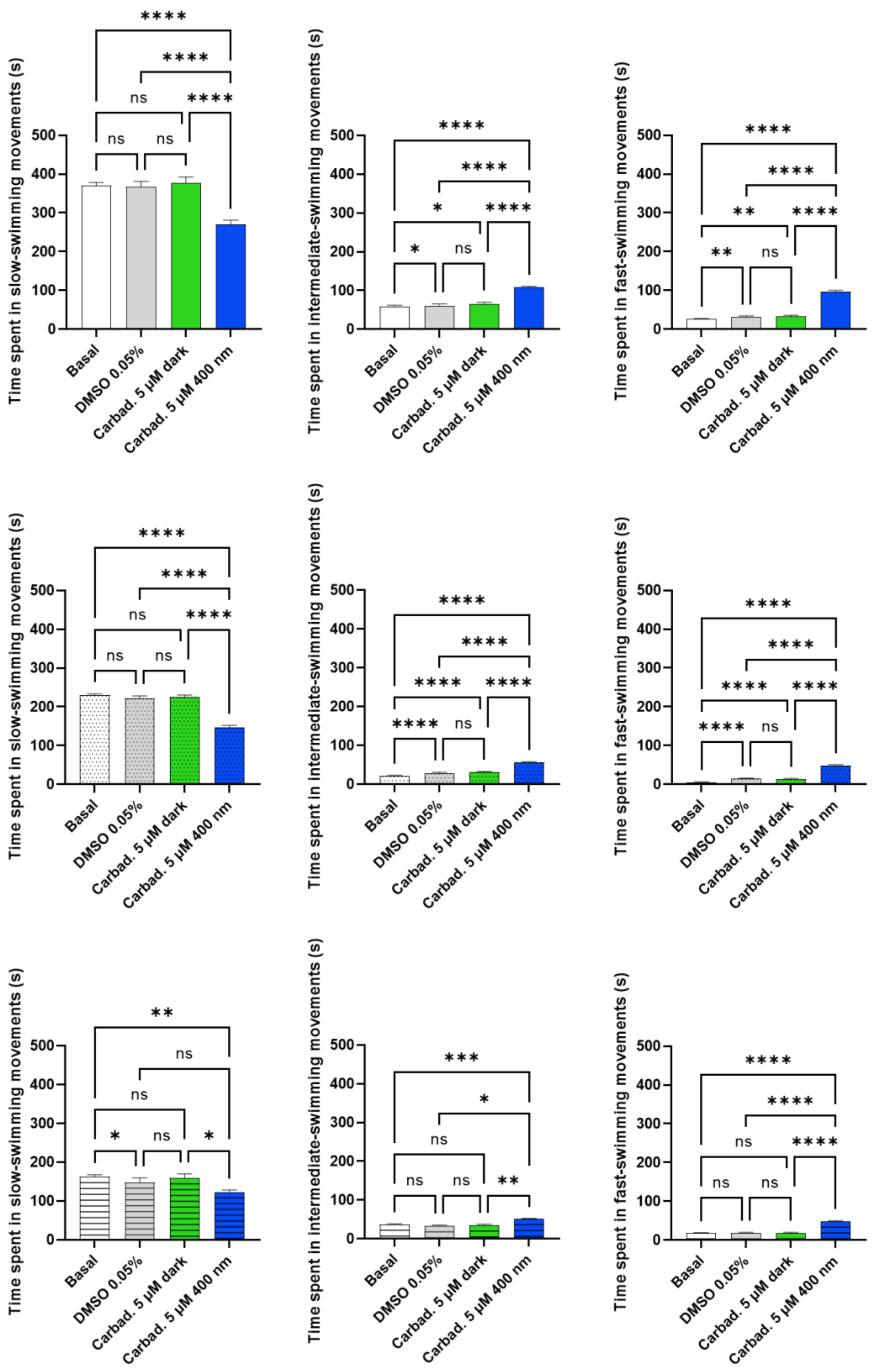

**Figure S15:** Time spent swimming at different speeds for larvae in basal condition, treated with vehicle (DMSO 0.05 %), benchtop Carbadiazocine and 400 nm pre-illuminated Carbadiazocine. Smooth bars represent the data for the complete experiment, dotted bars during the no-stimuli period and striped bars during the stimuli period. Data represented as mean ± S.E.M. ****p<0.0001; ***p<0.001; **p<0.01; *p<0.05; ns = not significant, Mixed-effect analysis followed by Tukey's multiple comparisons test.

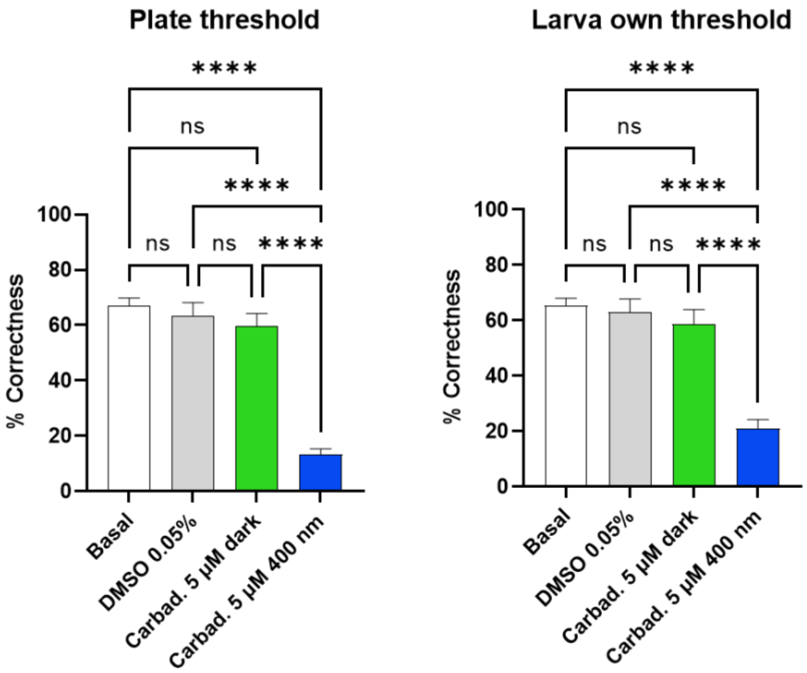

**Figure S16:** Correctness in the Optomotor response assay in free-swimming larvae setting different thresholds for correctness. The left panel shows the percentages of correctness setting as threshold for a positive response the average vertical distance travelled by the whole plate (ten fishes) during the no stimuli period 30 seconds bins in basal condition, while for the right panel the threshold was set as the average vertical distance travelled by each larva as its own control during the no stimuli period 30 seconds bins in basal condition. Data represented as mean ± S.E.M. ****p<0.0001; ***p<0.001; **p<0.01; *p<0.05; ns = not significant, Mixed-effect analysis followed by Tukey's multiple comparisons test.
